# NRF1 Association with AUTS2-Polycomb Mediates Specific Gene Activation in the Brain

**DOI:** 10.1101/2021.03.30.437620

**Authors:** Sanxiong Liu, Kimberly A Aldinger, Chi Vicky Cheng, Takae Kiyama, Mitali Dave, Hanna K. McNamara, Stefano G Caraffi, Ivan Ivanovski, Edoardo Errichiello, Christiane Zweier, Orsetta Zuffardi, Michael Schneider, Antigone S. Papavasiliou, M. Scott Perry, Megan T Cho, Astrid Weber, Andrew Swale, Tudor C. Badea, Chai-An Mao, Livia Garavelli, William B Dobyns, Danny Reinberg

## Abstract

The heterogeneous complexes comprising the family of Polycomb Repressive Complex 1 (PRC1) are instrumental to establishing facultative heterochromatin that is repressive to transcription. Yet, two PRC1 species, PRC1.3 and PRC1.5, are known to comprise novel components, AUTS2, P300, and CK2 that convert this repressive function to that of transcription activation. Here, we report that patients harboring mutations in the HX repeat domain of AUTS2 exhibit defects in AUTS2 and P300 interaction as well as a developmental disorder reflective of Rubinstein-Taybi syndrome, which is mostly associated with a heterozygous pathogenic variant in *CREBBP/EP300*. As well, the absence of AUTS2 gives rise to a mis-regulation of a subset of developmental genes and curtails motor neuron differentiation from embryonic stem cells in the context of a well-defined system. Moreover, the transcription factor, Nuclear Respiratory Factor 1 (NRF1) exhibits a novel and integral role in this aspect of the neurodevelopmental process, being required for PRC1.3 recruitment to chromatin.

## INTRODUCTION

The establishment, maintenance, inheritance and regulated dissolution of facultative heterochromatin(Trojer and Reinberg, 2007) is paramount to the developmental processes that give rise to the distinct cellular identities comprising multi-cellular organisms. The Polycomb group (PcG) of proteins are required for the formation and integrity of facultative heterochromatin as a function of the presence of intracellular signals that occur during development, as well as for the maintenance of adult tissue-specific gene expression profiles(Bonasio et al., 2010; Margueron and Reinberg, 2011; Di Croce and Helin, 2013). Indeed, our findings demonstrate that in contrast to chromatin domains engaged in active transcription, it is the repertoire of repressive chromatin domains that are conveyed to daughter cells upon DNA replication, thereby maintaining cellular identity(Escobar et al., 2019, 2021).

Two multi-subunit complexes, Polycomb Repressive Complex-1 and −2 (PRC1 and PRC2, respectively) comprise a defined subset of PcG proteins(Schuettengruber et al., 2017), and act in concert to establish facultative heterochromatin. PRC2 comprises the sole activity that catalyzes mono-, di, and tri-methylation of histone H3 at lysine 27 (H3K27me1, -me2, -me3, respectively), with chromatin domains comprising H3K27me2/me3 forming the platform for chromatin compaction(Lau et al., 2017; Oksuz et al., 2018; Yu et al., 2019; Kim and Kingston, 2020). PRC1 complexes comprise other subsets of the PcG protein family. We and others previously characterized at least six heterogeneous PRC1 subcomplexes, each of which comprise one of the six Polycomb Group Ring Finger (PCGF1-6) components and RING1A and/or RING1B(Gao et al., 2012; Tavares et al., 2012; Hauri et al., 2016). This heterogenicity of the PRC1 complexes resulted in their classification into two major PRC1 subcomplexes. Canonical PRC1 (cPRC1) comprise one of several CBX components that bind to chromatin containing nucleosomes decorated with H3K27me3 catalyzed by PRC2, thereby resulting in chromatin compaction and thus, transcription repression(Min et al., 2003; Francis et al., 2004; Gao et al., 2012; Lau et al., 2017; Kim and Kingston, 2020). The non-canonical set of PRC1 (ncPRC1) comprise either RYBP or YAF2 that stimulate the catalysis of H2A mono-ubiquitinated lysine 119 (H2AK119ub1) through the common PRC1 subunit, RING1A/B. Of note, incorporation of CBX or either RYBP or YAF2 into PRC1 is competitive such that cPRC1 is devoid of RYBP and YAF2 and ncPRC1 is devoid of CBX(Wang et al., 2010; Gao et al., 2012). The joint effect of PRC2/ncPRC1 arises from a PRC2 accessory partner, Jarid2, which stimulates PRC2 activity(Li et al., 2010; Pasini et al., 2010). Jarid2 reportedly interacts with the ncPRC1-mediated product, H2AK119ub1(Kasinath et al., 2021). Interestingly, while this joint recruitment of PRC2 with either cPRC1 or ncPRC1 is distinct, both of these versions of PRC1 are found in proximity at select genome-wide regions(Gao et al., 2012; Scelfo et al., 2019). Most importantly, this PRC2/PRC1 network is paramount to fostering the appropriate profiles of facultative heterochromatin evident during development and in adulthood(Margueron and Reinberg, 2011; Aloia et al., 2013; Schuettengruber et al., 2017).

Our previous characterization of PRC1 complexes also revealed that a subset of ncPRC1 that comprise either PCGF3 (ncPRC1.3) or PCGF5 (ncPRC1.5) unexpectedly contained three non-PcG proteins: AUTS2, CK2 and P300(Gao et al., 2012, 2014). Remarkably, we found that these proteins hijack the normally repressive ncPRC1 complex and convert it into a transcriptional activator(Gao et al., 2014). Indeed, CK2 mediates the phosphorylation of the RING1A/B subunit of PRC1, thereby thwarting its catalysis of H2AK119ub1. Moreover, AUTS2 interacts with PCGF3/-5 and most importantly, recruits P300/CBP, a known transcriptional co-activator possessing histone acetyltransferase activity(Bannister and Kouzarides, 1996; Ogryzko et al., 1996). The presence of AUTS2, P300/CBP and CK2 within two ncPRC1 complexes (PRC1.3/1.5) converts the function of these complexes into that of transcription activation, rather than repressive activity(Gao et al., 2014). These findings point to AUTS2 having a profound impact on gene expression in the context of defined aspects of development. Indeed, AUTS2-ncPRC1.3 is important during development of the central nervous system (CNS) as well as in the post-developmental stage (see below). In contrast, AUTS2-ncPRC1.5 appears to function in the establishment and maintenance of other lineages of differentiation (our unpublished results). Notably, ncPRC1.3/-1.5 comprises another non-PcG protein, FBRSL1, which shares a highly similar protein sequence with AUTS2(Gao et al., 2012). Interestingly, AUTS2 and FBRSL1 bind competitively to the PCGF subunit of ncPRC1.3/-1.5(Gao et al., 2014).

The gene encoding AUTS2 was designated as such based on the identification of its translocation in a pair of monozygotic twins that were diagnosed with autism(Sultana et al., 2002); yet its role in Autism Spectrum Disorders (ASD) is still putative. Nonetheless, the role of AUTS2 in neurodevelopment has been more widely established through the identification of variants in *AUTS2* that are associated with variable intellectual disability (ID), outgoing social behavior, mild microcephaly, and selected co-morbid features of autism that include obsessive-compulsive behavior(Beunders et al., 2013; Oksenberg and Ahituv, 2013; Hori and Hoshino, 2017). Various reports indicate the association of AUTS2 variants with other pronounced neurological diseases, including epilepsy, bipolar disorder, attention deficit hyperactive disorder, and alcohol dependency(Hattori et al., 2009; Mefford et al., 2010; Elia et al., 2010; Kapoor et al., 2013). Notably, a thorough and informative report analyzed AUTS2 binding to chromatin genome-wide in mouse E16.5 forebrain, finding that AUTS2 is bound to gene promoters and enhancers whose function appears to be important during neurodevelopment(Oksenberg et al., 2014), and pointing to its substantive role in activating genes required for appropriate CNS development and function.

Given that AUTS2 interacts with P300, converting ncPRC1.3 into a transcriptional activator, it is notable that human heterozygous pathogenic variants in *EP300* or *CREBBP* (CREB Binding Protein/CBP) are associated with Rubinstein-Taybi syndrome (RSTS), a neurodevelopmental disorder characterized by distinctive facial features, broad and angulated thumbs, short stature, and intellectual disability(Stevens, 1993; Ajmone et al., 2018). Here, we report heterozygous *de novo* variants in *AUTS2* in patients who exhibit a severe phenotype overlapping that of RSTS. Notably, these new AUTS2 variants are defective in P300/CBP interaction, underscoring the biological relevancy of P300/CBP incorporation into AUTS2-ncPRC1.3 with respect to appropriate neurodevelopment and brain function in human.

We further extended our studies to understand the means by which AUTS2-ncPRC1.3 accesses specific chromatin sites in the brain and identified the transcription factor, Nuclear Respiratory Factor 1 (NRF1), as being instrumental to this process. Previous studies implicated NRF1 in mitochondrial biogenesis(Scarpulla, 2011) and appropriate development of the retina(Hsiao et al., 2013; Kiyama et al., 2018). Here we demonstrate that NRF1 mediates AUTS2-ncPRC1.3 recruitment to a subset of neurodevelopmental genes during differentiation of mouse embryonic stem cells to motor neurons, as well as in the mouse brain during early development. These findings expose a novel and key role for NRF1 in facilitating appropriate AUTS2-ncPRC1.3-mediated activation of genes involved in neurodevelopment as a consequence of AUTS2-P300 interaction.

## RESULTS

### ncPRC1.3 occupies active genes during early development in the mouse brain

The *Auts2* gene encodes 2 major transcript isoforms in the mouse brain(Hori et al., 2014). The full-length (FL) mouse Auts2 transcript (Auts2-1, Figure 1A) includes 19 exons, and produces the long isoform of AUTS2 (AUTS2_L, 1-1261 aa, Figure 1B). Another Auts2 transcript (Auts2-2, Figure 1A) arises from a transcriptional start site near exon 7 and comprises a translational start site in the middle of exon 8 that gives rise to the short form of AUTS2 protein (AUTS2_S, 458-1261 aa, Figure 1B)(Hori et al., 2014). Both AUTS2 isoforms contain a PY motif and an HX repeat that in the case of AUTS2-L is located between two proline rich regions (PR1 and PR2) in the N-terminus (Figure 1B). The PY motif is a potential WW-domain-binding region present in various activating transcription factors(Sultana et al., 2002). The HX repeat (aa 525-542) comprises alternating HQ (x6) or HT (x3) residues, mutations of which in two other genes (*ATN1* and *RERE*), are related to neurodevelopmental disorders(Jordan et al., 2018; Palmer et al., 2019).

**Figure 1.**
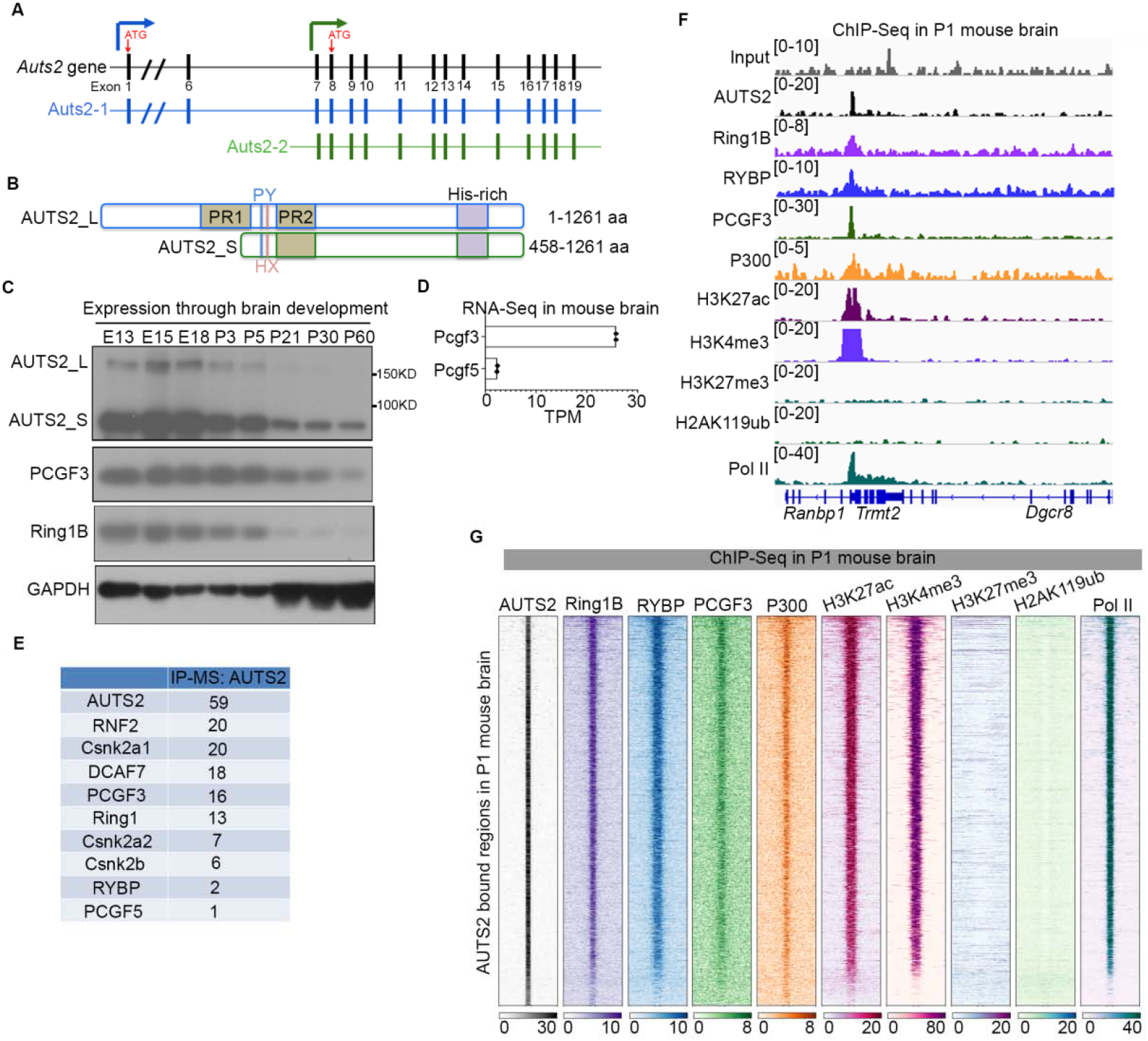
AUTS2-ncPRC1.3 targets active genes in mouse brain. (A) Schematic showing the mouse *Auts2* gene structure and its two major transcripts in the mouse brain. Red arrows indicate the translational start codon used for each transcript. (B) Schematic showing the domains of the long and short isoforms of mouse AUTS2 protein. PR, proline-rich region; PY, PPPY motif; HX, HX repeat motif, comprising alternating HQ (x6) or HT (x3) residues; His-rich, eight histidine repeats. (C) Expression of AUTS2 and core ncPRC1.3 components in the mouse brain. Immunoblotting was performed with whole brain extracts at various developmental stages, as indicated. (D) Bar graphs showing the value of transcripts per kilobase million (TPM) for Pcgf3 and Pcgf5 revealed by RNA-Seq performed from whole brain lysate at postnatal day 1. (E) Proteomic mass spectrometry results of immunoprecipitation (IP) using AUTS2 antibody in whole brain lysate at postnatal day 1. (F) IGV browser views showing ChIP-seq profile for input, AUTS2, RING1B, RYBP, PCGF3, P300, H3K27ac, H3K4me3, H3K27me3, H2AK119ub1 and RNA Polymerase II (PolII) at the representative loci. ChIP-seq was performed in whole brain lysate at postnatal day 1. (G) Heatmap showing AUTS2, RING1B, RYBP, PCGF3, P300, H3K27ac, H3K4me3, H3K27me3, H2AK119ub1 and RNA Polymerase II ChIP-seq signals centered on AUTS2 bound regions (±5 kb). ChIP-seq was performed in whole brain lysate at postnatal day 1.

To investigate the role of AUTS2 specifically in the context of its associated ncPRC1 complex in brain development, we examined its expression and that of the core PRC1 components in the mouse brain throughout early development. Western blot analysis in whole mouse brain lysates using an antibody against the AUTS2 C-terminus [1160–1259 amino acids of human AUTS2 protein(Gao et al., 2014)], showed expression of both FL AUTS2 protein (approximately 170 kDa) and its shorter form (approximately 95 kDa) with the latter being predominant (Figure 1C). The expression of both isoforms gradually decreased throughout early development (Figure 1C). Accordingly, both Ring1B and PCGF3 expression subsided dramatically from postnatal day 5 (P5) (Figure 1C).

As AUTS2 is incorporated into both ncPRC1.3 comprising PCGF3 and ncPRC1.5 comprising PCGF5 in 293 T-REx cells(Gao et al., 2012), we examined the expression of Pcgf3 and Pcgf5 from whole brain lysates at postnatal day 1. Interestingly, RNA-Seq data revealed that Pcgf3, but not Pcgf5, was predominantly expressed in the mouse brain (Figure 1D). Mass spectrometry (MS) analysis following co-immunoprecipitation (co-IP) experiments from brain lysate, using AUTS2 antibody recovered considerably more peptides from Pcgf3 than from Pcgf5 (Figure 1E). As previously reported(Gao et al., 2014), Two other components comprising ncPRC1.3, Ring1A/B and casein kinase 2 (CK2), were also observed (Figure 1E). Importantly, brain-specific conditional knockout of *Pcgf3* (*Pcgf3*^loxp/loxp^:*Nes*^Cre^) caused lethality (data not shown), suggesting a critical role for AUTS2-ncPRC1.3 during early brain development.

To better understand how AUTS2-ncPRC1.3 is involved in transcriptional regulation, we next characterized the genomic localization of AUTS2, P300, and the ncPRC1.3 components: RING1B, RYBP and PCGF3, by ChIP followed by deep sequencing (ChIP-seq) in whole brain lysate at postnatal day one. Consistent with our previously published ChIP-seq data in mouse brain, AUTS2 associated with ncPRC1.3 components including P300 in the promoter of active genes that were devoid of histone post-translational modifications (hPTMs) associated with transcription repression, and instead exhibited strong signals for hPTMs associated with active transcription, *e.g.* H3K27ac and H3K4me3 (Figure 1F). This finding was corroborated by a genome-wide analysis (Figure 1G), and together provide strong evidence that AUTS2-ncPRC1.3 is involved in active transcription in the mouse brain. GO analysis revealed that terms related to RNA processing and neuronal development were enriched in genes located in AUTS2-bound regions (Figure S1).

### Patients with mutations in the *AUTS2* HX repeat share features with Rubinstein-Taybi syndrome

We previously reported that a truncated form of AUTS2 protein (404 to 913 aa) was sufficient to mediate transcriptional activation through its recruitment of P300(Gao et al., 2014). Yet, the AUTS2 residues key to its interaction with P300 and the physiological relevance of AUTS2-P300 interaction during brain development remained largely unexplored. As part of an ongoing effort to identify genetic variants associated with developmental brain disorders(Aldinger et al., 2019), we detected a novel *de novo* missense variant in *AUTS2* in a boy with multiple congenital anomalies and a proposed diagnosis of RSTS. His phenotype was more severe than the syndrome previously reported among individuals with heterozygous *AUTS2* deletions(Beunders et al., 2013). To investigate the clinical phenotype associated with AUTS2 mutations, we identified 6 additional individuals with *de novo* intragenic variants that were clustered in exon 9 of the *AUTS2* coding region (NM_015570.2) (Figures 2A, S2 and Table S1). All 7 individuals displayed dysmorphic features and feeding difficulties in infancy, and most had moderate to severe intellectual disability and hypotonia (Table S1). Importantly, 5 of these individuals harbored mutations within a short histidine-rich HX repeat motif (aa 525-542) that comprises alternating HQ (x6) or HT (x3) residues, all of whom had severe phenotypes (Table S1). Our original proband (LR05-007) had a missense mutation, p.Thr534Pro, while the remaining four individuals (LR15-003, LR18-404, LR19-314, LR19-506) had an identical recurrent small deletion within exon 9: p.His535_Thr542del (Figure 2A). Notably, pathogenic variants in the HX repeat of two other genes [ATN1(Palmer et al., 2019) and RERE(Jordan et al., 2018)] are also associated with neurodevelopmental disorders (Figure 2B).

**Figure 2.**
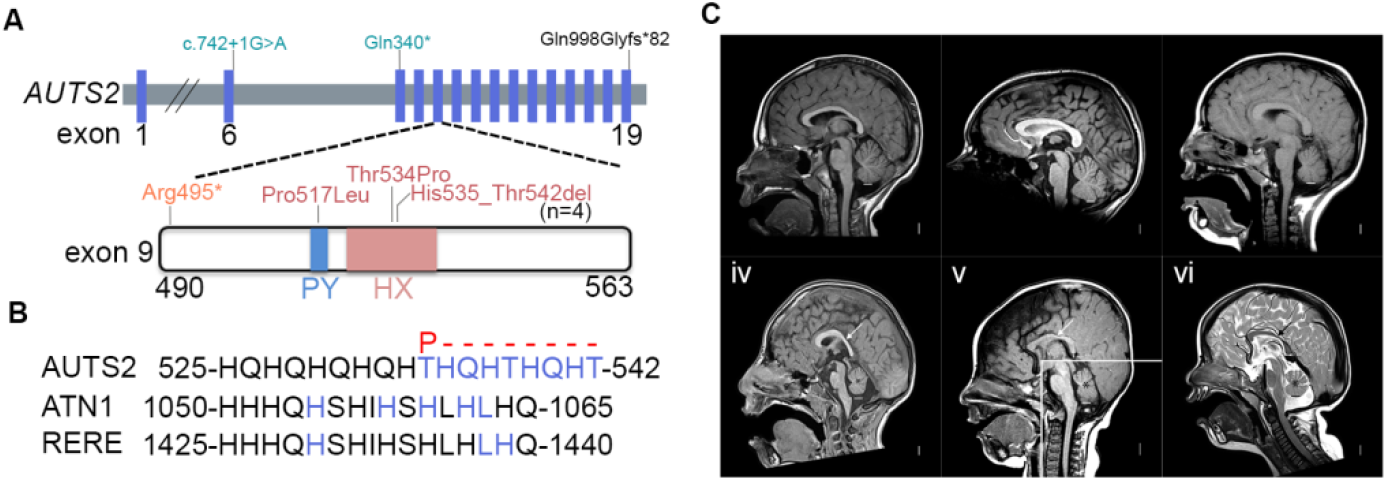
Patients with mutations in AUTS2 HX repeat have features that overlap with Rubinstein-Taybi syndrome. (A) Schematic illustrating mutations in the *AUTS2* gene from individual patients as identified through trio-based exome sequencing. Mutations resulting in similar clinical features are labeled with the same color. PY, PPPY motif; HX, HX repeat motif, comprising alternating HQ (x6) or HT (x3) residues. (B) Mutations in the HX repeat of AUTS2 compared with those reported for ATN1 and RERE. The mutated residues are labeled in blue. The variant substitutions for AUTS2 present in affected individuals are indicated above the sequence alignment (red). (C) Magnetic resonance images from 5 individuals with missense variants in AUTS2 exon 9 and a normal control. The three midline sagittal images in the top row all show normal midline structures. They come from a normal control (**i**) a girl with a missense variant (p.Thr534Pro) in the PY motif (**ii**) and a boy with the recurrent INDEL (p.His535_Thr542 del) in the HX motif (**iii**). The three midline sagittal iimages in the bottom row come from a boy with a missense mutation in the HX domain (p.Thr534Pro) and two boys with the recurrent INDEL in the HX motif. All three show thin show thin and dysplastic corpus callosa (arrows in **iv-vi**), and small cerebellar vermis (asterisks in **iv-vi**). The horizontal white or black lines mark the level of the obex, the usual lower extend of the vermis.

All 5 patients with mutations in the HX repeat of AUTS2 had a dysmorphic facial appearance dominated by features seen in RSTS, although less severe than classic RSTS (Table S1). RSTS is a complex multiple congenital anomaly syndrome characterized by short stature, distinctive facial features, and varying degrees of intellectual disability(RUBINSTEIN and TAYBI, 1963; Wiley et al., 2003). In most individuals, RSTS is associated with mutations in genes (*CREBBP, EP300*) encoding the CREB binding protein (CBP) or P300, or a microdeletion of 16p13.3 that includes CREBBP(Stevens, 1993). A clinical diagnosis of RSTS was suggested for 2 of the 5 individuals prior to genetic testing (Table S1). The two other patients with mutations outside of the HX repeat domain did not exhibit a phenotype overlapping that of RSTS (Table S1), but display other neurological defects. Moreover, neuroimaging studies from 4 of the 5 patients with mutations in the HX repeat showed hypoplasia of the corpus callosum (n=3), cerebellar hypoplasia and small posterior fossa (n=4), and Chiari malformation type 1 (n=1) (Figure 2C and Table S1). Such brain malformations are also reported in individuals with classic RSTS(Cantani and Gagliesi, 1998; Ajmone et al., 2018). These findings led us to hypothesize that the HX repeat in AUTS2 coordinates with CBP/P300 in regulating proper gene expression in the brain.

### The HX repeat in AUTS2 interacts with P300, crucial for activating transcription

Given that mutations in both the AUTS2 HX repeat and CBP/P300 are associated with RSTS, we next sought to determine whether the RSTS phenotype observed could be due to disrupted interactions between AUTS2 and CBP/P300. To this end, we initially expressed Flag-tagged AUTS2, either wild-type (WT) or patient-derived mutant forms including two variants within the HX repeat (T534P and 535-542aa del) and one outside this region (P517L within the PY motif) in 293 T-REx cells (Figures 3A and 2A). Strikingly, co-IP experiments using Flag-WT or Flag-mutant AUTS2 revealed that both mutations within the HX repeat (T534P and 535-542aa del), but not P517L, disrupted interaction with P300 (Figure 3B). This finding is in accordance with the clinical diagnosis of RSTS for patients harboring mutations within the HX repeat, but not the P517L mutation (Table S1). Of note, WT and all AUTS2 variants interacted stably with Ring1B (Figure 3B), suggesting that AUTS2 regions outside the PY motif and HX repeat could mediate AUTS2 incorporation into the ncPRC1 complex. Importantly, reciprocal co-IP experiments performed against endogenous P300 confirmed that its interaction with AUTS2 was disrupted when AUTS2 was mutant in its HX repeat (Figure 3C). Similar results were observed using a more relevant system: cells undergoing differentiation into motor neurons (see below).

**Figure 3.**
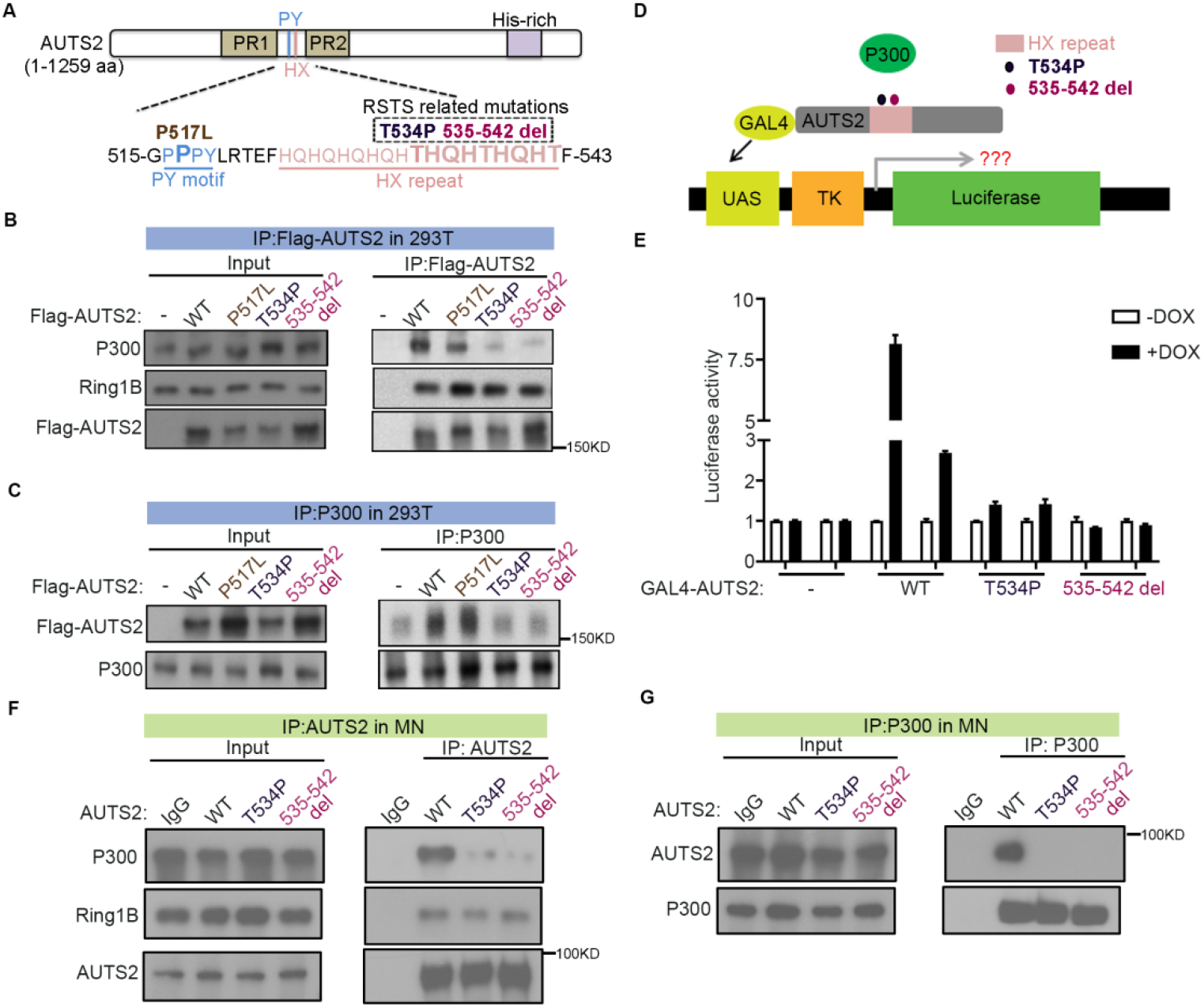
Mutations in AUTS2 HX repeats disrupt both P300 binding and AUTS2-mediated transcription activation. (A) Schematic showing human AUTS2 variants constructed and expressed in 293 T-REx cells. The two variants within the HX repeat are associated with Rubinstein-Taybi syndrome. (B-C) Western blots show co-IP results from nuclear extract of 293 T-REx cells expressing Flag– AUTS2, either WT or mutant versions as indicated, using Flag antibody (B) or reciprocal IP using P300 antibody (C). (D) Schematic of the reporter construct for the luciferase assay in the context of GAL4-AUTS2, either WT or mutant versions as indicated. (E) Luciferase activity in cells expressing GAL4-AUTS2, either WT or mutant versions before and after doxycycline treatment. (F-G) Western blots show co-IP results from nuclear extract of motor neuron differentiated from WT and *Auts2* HX mutant (T534P and 535-542 aa deletion respectively) mESC as indicated, using AUTS2 antibody (F) or reciprocal IP using P300 antibody (G).

Considering that P300 is required for AUTS2-mediated transcriptional activation(Gao et al., 2014), we next wondered whether AUTS2 mutants in the HX repeat lose the ability to activate transcription due to their disrupted interaction with P300. To this end, GAL4–AUTS2, either WT or mutant in the HX repeat, or GAL4 alone were inducibly expressed in 293 T-REx cells containing an integrated luciferase reporter with a UAS comprising five consecutive GAL4 DNA binding sites (Figure 3D). Importantly, we found a severe defect in doxycycline-mediated induction of luciferase activity in the case of the GAL4-AUTS2 mutants compared to WT (Figure 3E).

Finally, to confirm the critical role of the HX repeat domain in mediating interaction between AUTS2 and P300 in a more relevant system (*in vitro* neuronal differentiation), we generated mouse embryonic stem cells (mESC) genetically modified to harbor one of two mutations in the AUTS2 HX repeat domain at the endogenous locus (T534P or a 535-542 aa deletion, Figures S3A and S3B, respectively), which we could then differentiate toward the neuronal lineage (see Figure 5A for more details). Introduction of either mutation at the endogenous locus did not change the expression level of AUTS2 (Figure S3C). Consistent with our observations in 293T cells, patient-derived mutations in the AUTS2 HX repeat domain disrupted its interaction with P300 but not its incorporation into the ncPRC1 complex (Figures 3F and 3G). Altogether, these results establish that the AUTS2 HX repeat (aa 525–542) engages in P300 interaction, pointing to the critical role of this minimal region in AUTS2-mediated transcriptional activation and corroborating the consequences of the HX mutation in Rubinstein-Taybi syndrome (RSTS).

### AUTS2 and NRF1 co-localize within chromatin and interact in the mouse brain

To identify the factor(s) involved in the key process by which AUTS2 is recruited to chromatin, we first determined the motifs of transcription factors (TFs) enriched in AUTS2-bound sites in the mouse brain and identified an overrepresentation for that of Nuclear Respiratory Factor 1 (NRF1) (Figure 4A). NRF1 is a TF known for its role in mitochondrial biogenesis(Scarpulla, 2011), and binds to GC-rich DNA elements in promoters of many mitochondrial biogenesis-related genes(Evans and Scarpulla, 1990; Gleyzer et al., 2005). As well, the protein is associated with the regulation of neurite outgrowth(Chang et al., 2005; Tong et al., 2013) and exhibits essential roles in retinal development(Hsiao et al., 2013; Kiyama et al., 2018), yet its function and regulation in the CNS is largely unknown. To validate our computational prediction, we performed ChIP-seq for NRF1 using 2 different antibodies and lysates from whole mouse brain isolated at postnatal day one and ascertained that the majority of AUTS2 peaks (1545 of 2005 peaks in total) were associated with genomic sites bound by NRF1 (Figures 4B and 4C).

**Figure 4.**
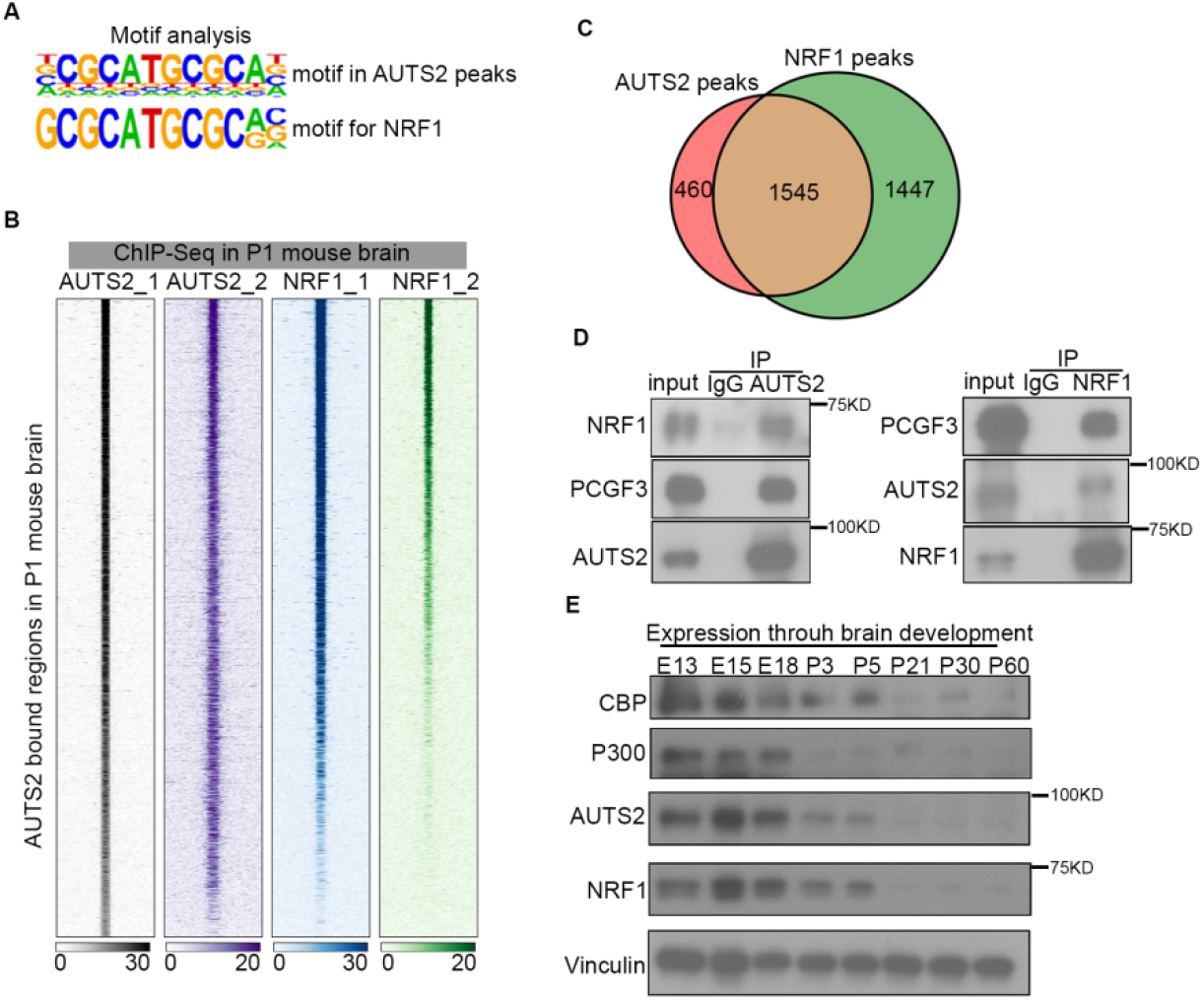
Transcription factor NRF1 colocalizes with AUTS2 on chromatin and physically interacts with AUTS2 in mouse brain. (A) Top, motif analysis of AUTS2-bound regions in mouse brain using HOMER. Bottom, known motif for transcription factor NRF1. (B) Heatmap showing AUTS2 and NRF1 ChIP-seq signals centered on AUTS2-bound regions (±5 kb) with two replicates. ChIP-seq was performed in whole brain lysate at postnatal day 1. (C) Venn diagram showing the extent of overlap for AUTS2- and NRF1-bound regions revealed by ChIP-seq performed in whole brain lysate at postnatal day 1. (D) Reciprocal co-immunoprecipitation and western blot analyses of AUTS2 and NRF1 interaction in whole brain lysate at postnatal day 1. (E) Expression of CBP, P300, AUTS2 and NRF1 in mouse brain. Immunoblotting was performed with whole brain extracts at various developmental stages, as indicated.

**Figure 5.**
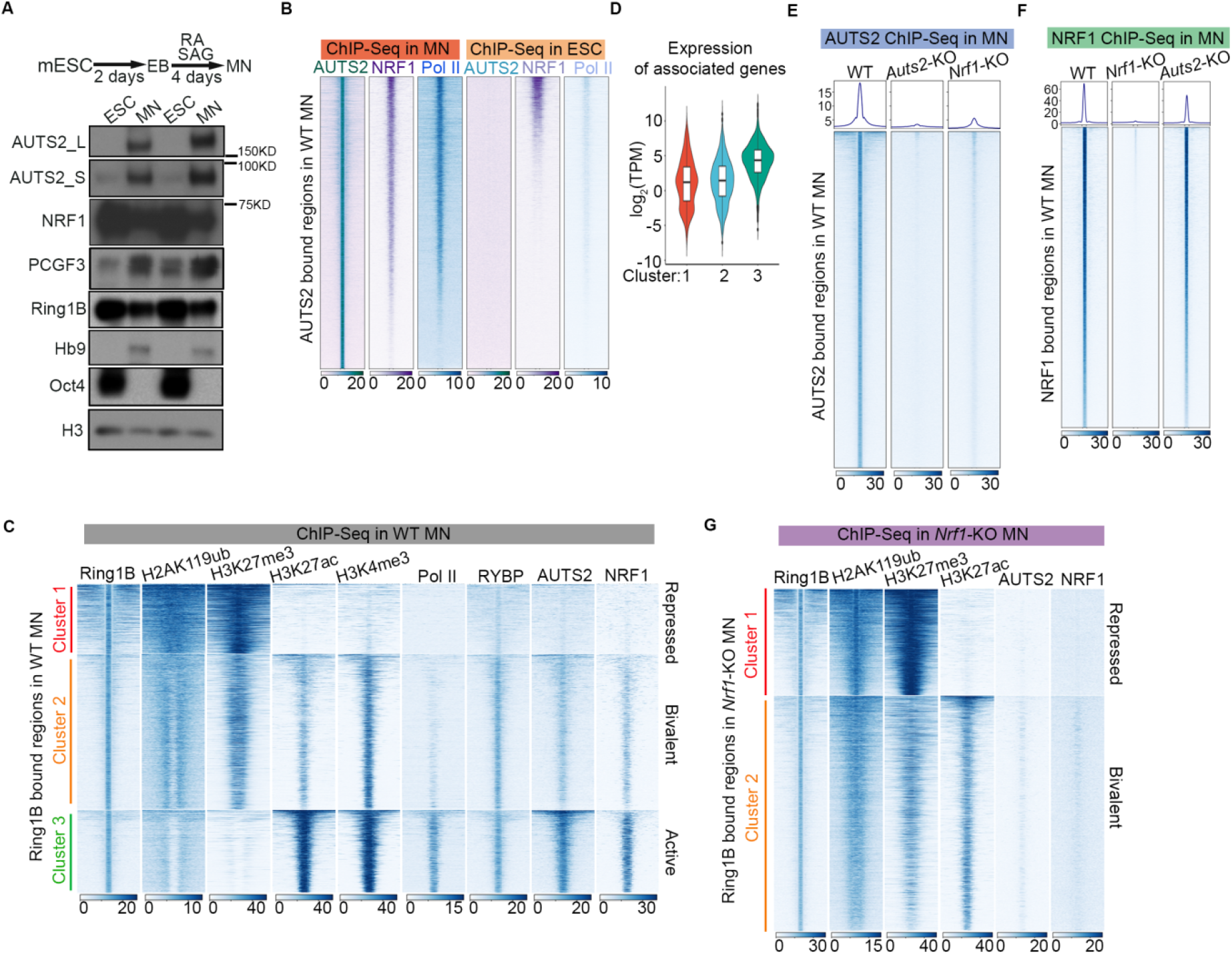
NRF1 is crucial for AUTS2-ncPRC1.3 associated active transcription in MN. (A) The schematic at top depicts the protocol for differentiation of mouse embryonic stem cells (mESC) to motor neurons (MN) using retinoic acid (RA) and smoothened agonist (SAG). EB, embryoid bodies. Below is a western blot showing the expression of AUTS2, NRF1, PCGF3 and Ring1B in ESC and MN. (B) Heatmap showing AUTS2, NRF1 and RNA Pol II ChIP-seq signals centered on AUTS2 bound regions identified in WT MN (±5 kb). ChIP-seq was performed in ESC and MN. (C) k-means clustering of RING1B, H2AK119ub, H3K27me3,H3K27ac, H3K4me3, RNA Pol II, RYBP, AUTS2 and NRF1 ChIP-seq signals from WT MN centered on Ring-B bound regions identified in WT MN (±5 kb). (D) Violin plot of the log_2_(TPM) of the genes assigned to each cluster [as indicated in (C)], quantified from RNA-seq in WT MN. (E) Average density profiles (top) and heatmap (bottom) showing AUTS2 ChIP-seq signals from WT, *Auts2*-KO and *Nrf1*-KO MN centered on AUTS2-bound regions identified in WT MN (±5 kb). (F) Average density profiles (top) and heatmap (bottom) showing NRF1 ChIP-seq signals from WT, *Nrf1*-KO and *Auts2*-KO MN centered on NRF1-bound regions identified in WT MN (±5 kb). (G) k-means clustering of RING1B, H2AK119ub, H3K27me3, H3K27ac, AUTS2 and NRF1 ChIP-seq signals from *Nrf1*-KO MN centered on Ring1B-bound regions identified in *Nrf1*-KO MN (±5 kb).

Based on this high overlap between chromatin-bound AUTS2 and NRF1, we next tested the possibility that NRF1 might physically interact with AUTS2, thereby contributing to their co-localization. Reciprocal co-IP assays using endogenous proteins revealed that indeed, NRF1 physically associates with AUTS2 in the mouse brain (Figure 4D). Notably, the core component of AUTS2-ncPRC1.3, PCGF3, also co-immunoprecipitated with NRF1 indicating that AUTS2 interacts with NRF1 within the context of the ncPRC1.3 complex in mouse brain (Figure 4D). Moreover, the expression of NRF1 recapitulated the pattern of AUTS2-ncPRC1.3 and CBP/P300 expression during early brain development (Figures 4E and 1C). These data strongly suggest that NRF1 contributes to the recruitment of AUTS2 and its associated ncPRC1 complex, although it is also clear that both have independent targets, likely due to their additional functions and partnership with other factors.

### AUTS2 and NRF1 colocalize with ncPRC1.3 at actively transcribed loci in cells induced to differentiate to motor neurons

To understand the underlying mechanism that coordinates both the chromatin binding of and the regulation of transcription by AUTS2-ncPRC1.3 and NRF1, we utilized the system by which differentiated motor neurons (MN) are attained *in vitro*(Wichterle et al., 2002; Mazzoni et al., 2013; Narendra et al., 2015). Under these conditions, the expression of both AUTS2 and PCGF3 was significantly up-regulated in MN, while that of NRF1 decreased to approximately half of its expression in mESC at both the protein (Figure 5A) and RNA levels (Figure S4A). In contrast, the overall level of PRC1 complex as reflected by that of RING1B was down-regulated (Figures 5A and S4A), consistent with its essential role in maintaining mESC identity(Endoh et al., 2008). To complement our previously published ChIP-seq data for RNA polymerase II (RNAPII) in both mESC and MN(Narendra et al., 2015; LeRoy et al., 2019), we performed similar ChIP-seq for AUTS2 and NRF1. The majority of regions that gained AUTS2 binding upon MN differentiation, also accumulated NRF1 binding (Figure 5B). Importantly, these regions became actively transcribed during differentiation as evidenced by an increase in RNAPII binding (Figure 5B).

To examine whether AUTS2 cooperates with ncPRC1.3 for active transcription in MN, we next analyzed the genome-wide distribution of a set of hPTMs (H2AK119ub, H3K27me3, H3K27ac and H3K4me3), and the core PRC1 subunits, RING1B and RYBP. Consistent with previous studies in other systems(Kloet et al., 2016; Cohen et al., 2018; Loubiere et al., 2020), k-means clustering revealed three discrete classes of Ring1B-bound regions in MN (Figure 5C). In cluster 1, we observed strong and broad ChIP-seq signals for RING1B, H2AK119ub, and H3K27me3, and the absence of signals for H3K27ac, H3K4me3, RNAPII and the noncanonical PRC1 component, RYBP/YAF2 (RYBP antibodies do not distinguish between RYBP and YAF2), suggesting that these regions are co-repressed by PRC2 and canonical PRC1. The second cluster exhibited lower levels of H2AK119ub and H3K27me3, and increased levels of H3K27ac and H3K4me3, these last two mostly abundant at the peak center, suggesting that these Ring1B-bound regions featured bivalency(Bernstein et al., 2006; Voigt et al., 2013). The third cluster exhibited both H3K27ac- and H3K4me3-marked active regions enriched for developmental GO terms (Figure S4B). Surprisingly, cluster 3 exhibited elevated RYBP/YAF2 levels. Importantly, both AUTS2 and NRF1 binding were specifically enriched in cluster 3 (Figure 5C). Accordingly, genes flanking cluster 3 regions were expressed at levels significantly higher than those in clusters 1 and 2 (Figure 5D). These data strongly suggest that the transcription factor NRF1 associates with AUTS2-ncPRC1 to facilitate active transcription in motor neurons.

### NRF1 directs AUTS2-ncPRC1 chromatin binding

To ascertain whether AUTS2 binding to chromatin is dependent upon NRF1 binding or vice versa, we first performed ChIP-seq analysis for the presence of AUTS2 or NRF1 as a function of depleting either NRF1 or AUTS2, respectively. Exon 9 of the *Auts2* gene was targeted by CRISPR-Cas9 to remove both the long and short forms of the protein (Figures S4C-E, Table S2) and exon 4 of the *Nrf1* gene was targeted by CRISPR-Cas9 to remove NRF1 protein (Figures S4F-H, Table S2) in mESC from which MN were then derived. As a consequence of NRF1 depletion, most AUTS2-binding events were decreased in MN (Figures 5E and S5A). However, NRF1 ChIP-seq signals remained largely unaltered upon AUTS2 depletion (Figures 5F and S5B), demonstrating that binding of AUTS2 is NRF1-dependent, but NRF1 binding to chromatin is independent of AUTS2.

We next probed how NRF1-directed AUTS2 binding might modulate ncPRC1.3-associated active transcription. Under NRF1 depleted conditions, we performed ChIP-seq for Ring1B and a set of hPTMs (H2AK119ub, H3K27me3 and H3K27ac) followed by k-means clustering analysis. Remarkably, we did not recover the cluster of Ring1B-bound active regions (labeled by H3K27ac, but not by H3K27me3 or H2AK119ub), relative to the control (Figures 5G and 5C), implying that ncPRC1.3 was no longer associated with active transcription in the absence of NRF1. Collectively, these data demonstrate a pivotal role for NRF1 in facilitating ncPRC1.3-associated active transcription by directing AUTS2 binding to chromatin in motor neurons.

### The absence of AUTS2 or NRF1 leads to a defect in PNP to MN differentiation

Despite much evidence indicating that mutations in the *AUTS2* gene are associated with multiple neurodevelopmental disorders, including ASD(Oksenberg and Ahituv, 2013), and the essential role(s) of NRF1 in retinal development(Hsiao et al., 2013; Kiyama et al., 2018), that AUTS2 and NRF1 might coordinately regulate the process of neuronal differentiation was not previously recognized. To characterize gene expression changes during the transition from multipotent, posterior neural progenitors (PNPs) to terminally differentiated MNs upon ablation of AUTS2 or NRF1, we performed single-cell RNA sequencing (scRNA-seq) on MN differentiated from mESC either WT, or having a knockout (KO) of *Auts2* (*Auts2*-KO) or *Nrf1* (*Nrf1*-KO) (see Methods). We chose the newly developed Smart-seq3 technique(Hagemann-Jensen, 2020; Hagemann-Jensen et al., 2020), an improved version of Smart-seq2 with a 5’-unique molecular identifier RNA counting strategy and a much higher sensitivity that detects thousands more transcripts per cell. We obtained 632 high-quality (cells with >3000 detected genes) single-cell transcriptomes from all samples (WT, 228 cells; *Auts2*-KO, 221 cells; *Nrf1*-KO, 183 cells) for in-depth analyses (Figures 6A and S6A). To identify major cell types, we performed unsupervised clustering on a graph-based representation of the cellular gene expression profiles and 5 major clusters were visualized in a uniform manifold approximation and projection (UMAP) embedding(Butler et al., 2018; Becht et al., 2019) and represented by color-coded dashed-line circles (Figure 6A). Clusters were annotated according to known markers and previously established lineage information(Wichterle et al., 2002; Briggs et al., 2017), *i.e.*, posterior neural progenitor (PNP) expressing Sox3, posterior and ventral neural progenitor (PVNP) expressing Hoxd4, newborn motor neuron (NMN) expressing Neurog2, and motor neuron (MN) expressing Mnx1 and Chat (Figures S6B and S6C). Importantly, cells from all samples were clustered by cell type identity rather than sample identity (Figure 6A), indicating little or no batch effect. Notably, Auts2 and Nrf1 were highly expressed in all cell types (Figure S6C), suggesting their involvement in all of the different stages of differentiation.

**Figure 6.**
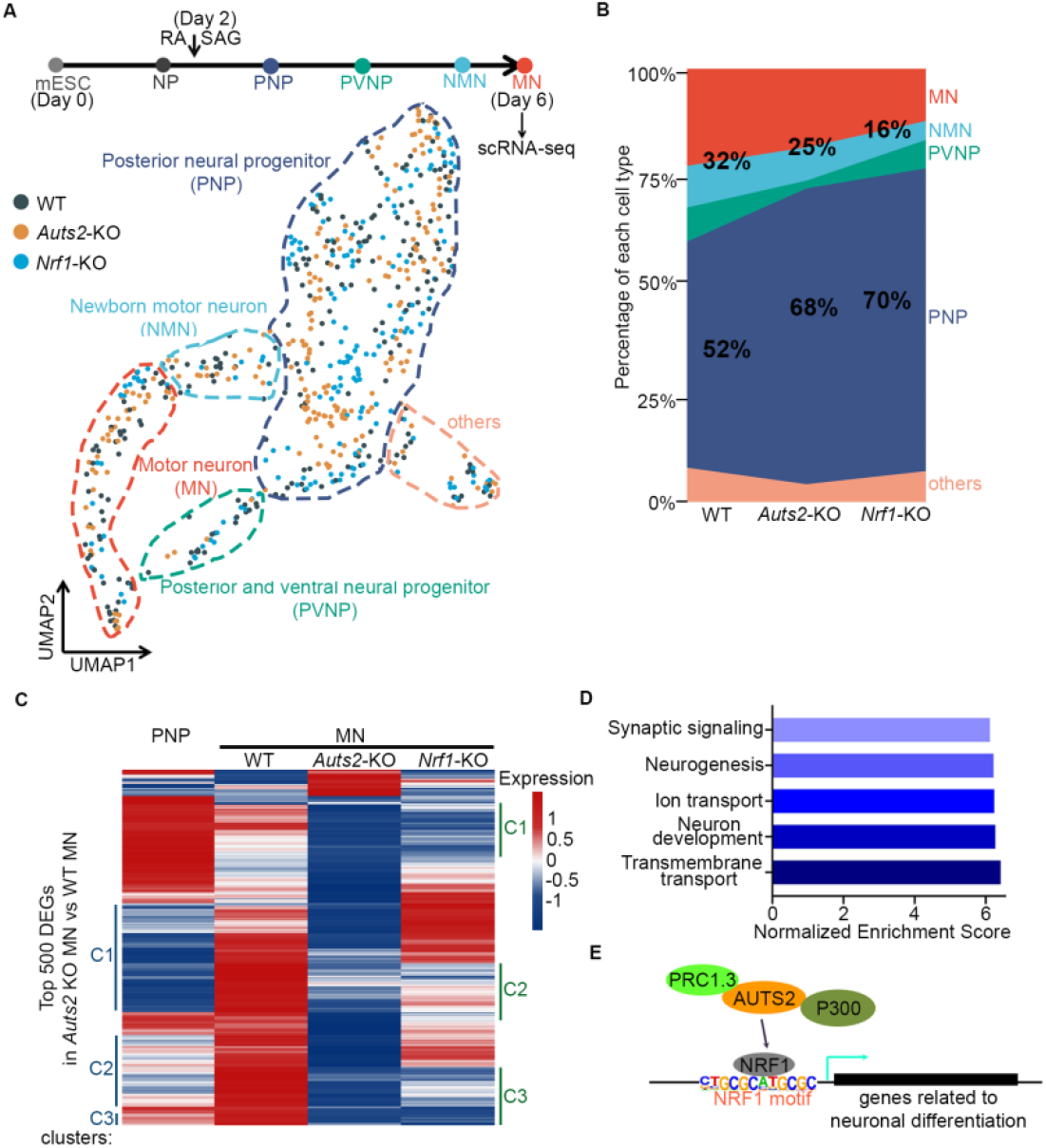
Defect in posterior neural progenitor differentiation into motor neuron in the absence of AUTS2 or NRF1 by scRNA-Seq. (A) The schematic at top depicts cell lineage transitions from mouse embryonic stem cells (mESC) to motor neurons (MN). MN (day 6) differentiated from WT, *Auts2* knockout and *Nrf1* knockout ESC were harvested for scRNA-Seq. Below is the dimensionality reduction (UMAP) of 632 cells from WT, *Auts2*-KO and *Nrf1*-KO samples, sequenced with the Smart-seq3 technique and colored by sample identity (WT, 228 cells; *Auts2*-KO, 221 cells; *Nrf1*-KO, 183 cells). Five cell classes revealed by unsupervised clustering of cellular transcriptomics are represented by color-coded circles of dashed-lines and annotated based on marker gene expression. Colors of dashed-lines match those for the cell types shown at top. (B) Proportional stacked area graph showing the abundances of each cell type in MN differentiated from WT, *Auts2*-KO and *Nrf1*-KO ESC. The percentage of motor neuron (MN+NMN) and posterior neural progenitor (PNP) in each sample are labelled. (C) Top 500 differentially expressed genes (DEGs) were identified by comparing *Auts2*-KO MN versus WT MN. Expression of these DEGs across WT PNP, WT MN, *Auts2*-KO MN and *Nrf1*-KO MN is shown by heatmap. Color scale represents the averaged and scaled expression values from each cell population. Labelling of clusters (C1, C2, C3) on left is based on gene expression differences in WT PNP, WT MN and Auts2-KO MN. Labelling of clusters (C1, C2, C3) on right is based on gene expression differences in WT MN, *Auts2*-KO MN and *Nrf1*-KO MN. (D) Bar plot summarizing results of gene set enrichment analysis for genes downregulated in *Auts2* KO MN versus WT MN. (E) Model depicting transcription factor NRF1-mediated recruitment of AUTS2-ncPRC1.3 resulting in activation of genes related to neuronal differentiation.

To pursue the potential functional requirement of AUTS2 and NRF1 for proper MN differentiation, we first compared the percentage of MN (including NMN and MN) and PNP under WT, *Auts2*-KO and *Nrf1*-KO conditions. The percentage of PNP was retained at a much higher level in both KO conditions (52% in WT, 68% and 70% in *Auts2*-KO and *Nrf1*-KO, respectively, Figure 6B). Moreover, a slight decrease in the percentage of terminally differentiated MN in *Auts2*-KO (32% to 25%) and a more severe defect in *Nrf1*-KO (32% to 16%) were observed (Figure 6B), indicating that *Auts2*-KO and *Nrf1*-KO result in a defect in PNP differentiation into MN. To gain more insight into the molecular mechanism by which AUTS2 and NRF1 contribute to the transition from PNP to MN, we further analyzed the differentially expressed genes (DEGs) specifically in the MN population from either WT or *Auts2*-KO (see Methods). Among the top 500 DEGs identified from WT MN and *Auts2*-KO MN, 458 genes were down-regulated in *Auts2*-KO, in accordance with the role of AUTS2 in transcriptional activation (Figures 6C, 1G and 5B)(Gao et al., 2014). Furthermore, about half of these 458 genes (205 of 458) were also down-regulated in *Nrf1*-KO MN compared to WT MN (C1, C2 and C3 labeled on the right, Figure 6C). To test whether NRF1-directed AUTS2 binding is required for the transcriptional activation of AUTS2-ncPRC1-associated active genes (cluster3 region, Figures 5C and 5G), we compared the differentially expressed genes in *Nrf1*-KO MN with the genes located in cluster3 and cluster1 regions from Ring1B ChIP-Seq. Consistent with the loss of AUTS2-ncPRC1 binding to active genes upon NRF1 knockout (Figures 5C and 5G), genes located in cluster3 regions were enriched in the down-regulated category instead of being up-regulated in *Nrf1*-KO MN (Figure S7). In contrast, very few of the genes located in cluster1 region (co-repressed by cPRC1 and PRC2) are affected in *Nrf1*-KO MN (Figure S7). These results strongly suggest that AUTS2 and NRF1 coordinately function in regulating transcriptional activation (Figures 5C and 5G).

We next asked whether the down-regulated genes in *Auts2*-KO MN reflected the WT transition from PNP to MN. Indeed, 186 of the 458 genes were normally up-regulated during PNP to MN differentiation in WT, but were defective in activation in *Auts2*-KO (C1, C2 and C3 labeled on the left, Figure 6C). For example, up-regulation of *Asic2*, a member of the sodium channel superfamily that regulates synaptic function(Zha et al., 2009) and of *Pnpla6*, a phospholipase that functions in neurite outgrowth(Guerreiro et al., 2015) were significantly attenuated under conditions of AUTS2 depletion. Finally, genes down-regulated in Auts2-KO MN were enriched for GO terms related to neuronal differentiation and function (Figure 6D). These data demonstrate that AUTS2 and NRF1 function coordinately to foster the appropriate differentiation of posterior neural progenitors to motor neurons by directly binding to and activating a subset of the relevant genes (Figure 6E).

## DISCUSSION

The results herein point to an instrumental role for the transcription factor NRF1 in facilitating chromatin access to the ncPRC1.3 comprising AUTS2, P300, CK2, PCGF3, and RYBP or YAF2. The function of this particular ncPRC1 has been converted from that of the typical PRC1 in facilitating transcription repression to that of a transcriptional activator(Gao et al., 2014). This conversion involves AUTS2-mediated recruitment of the P300 transcriptional co-activator and CK2-mediated phosphorylation of serine 168 of the integral PRC1 subunit, RING1, thereby thwarting PRC1-mediated monoubiquitination of H2AK119(Gao et al., 2014). Given that RYBP/YAF2 within other ncPRC1 complexes stimulate such RING1A/RING1B-mediated ubiquitination, their presence within ncPRC1.3 may indicate additional RYBP/YAF2 function(s). Importantly, as shown here, most of the genomic sites occupied by AUTS2 require NRF1, while most NRF1 sites are AUTS2-independent.

We noticed that the motif of transcription factor NRF1 is also significantly enriched in AUTS2-bound regions in a previous report(Oksenberg et al., 2014), although the enrichment is less dramatic than observed here. A recent study identified transcription factor USF1/2 as being key to PCGF3 chromatin binding in mESC(Scelfo et al., 2019), yet the motif corresponding to the DNA binding site of USF1/2 was not recovered in our study, suggesting that cell type/tissue specific mechanisms might dictate ncPRC1.3 recruitment to chromatin.

Instead, our findings direct attention to NRF1 in facilitating chromatin access by AUTS2, which is pivotal to the role of AUTS2 at a subset of genes involved in neurodevelopment. NRF1 has previously been shown to exhibit dimerization and to be subject to phosphorylation at several serine residues in its amino-terminus(Gugneja and Scarpulla, 1997). These phosphorylation events do not regulate NRF1 dimerization, but instead mutation of these sites compromise NRF1 DNA binding activity(Gugneja and Scarpulla, 1997). This report also indicates that CK2 could stimulate the DNA binding activity of NRF1 *in vitro*(Gugneja and Scarpulla, 1997). As CK2 is an integral component of ncPRC1.3 and inhibits its repressive activity by phosphorylating its RING1A/B component(Gao et al., 2014), the presence of CK2 might also foster NRF1 activity to promote ncPRC1.3-mediated transcription activation through AUTS2 interaction with P300. We speculate that CK2 exerts such a coordinated function, resulting in the optimal activation of ncPRC1.3-AUTS2 target genes in the brain.

Evidence involving NRF1 have highlighted its importance in mitochondrial biogenesis(Scarpulla, 2011), as well as in retinal development(Hsiao et al., 2013; Kiyama et al., 2018). Intriguingly, the pathways fostering mitochondrial integrity might be critical to those regulating distinct developmental pathways. While little is known about its role in development of the CNS, NRF1 is widely expressed within the CNS as evidenced by the *Nrf1^LacZ^* mouse line (Figure S8B). We attempted to delete NRF1 in the mouse brain from the embryonic stage in order to study NRF1-mediated AUTS2 recruitment, but such embryos did not survive, consistent with a previous report that *Nrf1*-null mouse embryos die between embryonic day 3.5 (E3.5) and E6.5(Huo and Scarpulla, 2001). Instead, we chose the *Tbr1^CreERT2^* line to strategically delete NRF1 in the adult mouse brain to examine the role of NRF1 in the CNS (Figures S8A-C). Importantly, we noticed several histological anomalies in these mutant mice (*Tbr1^CreERT2/+^: Nrf1^fx/fx^: Pou4f1^CKO/+^*), compared to control mice (*Tbr1^CreERT2/+^: Nrf1^fx/+^: Pou4f1^CKO/+^*), including a reduction in the size of the hippocampus and the width of the corpus callosum, as well as an enlarged lateral ventricle, indicating neuronal loss in both the cortex and hippocampus (Figures S8D and S8E). As well, a significant loss in retinal ganglion cells (RGCs) is observed in retinas collected from these mutant mice (Figures S8F and S8G).

It is important to emphasize that the reduced volume of corpus callosum observed in NRF1 mutant (*Tbr1^CreERT2/+^: Nrf1^fx/fx^: Pou4f1^CKO/+^*) mice is a key syndromic feature observed in RSTS patients(Cantani and Gagliesi, 1998), as well as in patients harboring mutations in the AUTS2 HX repeat domain as reported here, strongly supporting that NRF1 and AUTS2 function coordinately in regulating brain development. The corpus callosum (CC) connects the cerebral hemispheres and is the largest fiber tract in the brain(Edwards et al., 2014). During development, defects in neurogenesis, telencephalic midline patterning, neuronal migration and specification, axon guidance and post-guidance development can interrupt CC formation(Reyes et al., 2020).

Our findings also point to an instrumental role for the AUTS2 HX repeat domain given its requirement for AUTS2 interaction with P300/CBP. Importantly and as shown here, AUTS2 variants in this domain exhibit profound clinical and transcriptional effects *in vivo*. Given the existence of ncPRC1.3 in which AUTS2 conveys transcription activation, along with the reported association of AUTS2 haploinsufficiency in AUTS2-syndrome(Beunders et al., 2013), and possibly in Autism Spectrum Disorders(Sultana et al., 2002), AUTS2 had appeared key for modulating appropriate neurodevelopment. Here, we identified a critical role for its HX repeat domain in mediating AUTS2 interaction with P300 and resultant transcriptional activation by interrogating mutations found in individuals exhibiting a distinct and severe neurodevelopmental syndrome that overlaps with RSTS. Our *in vitro* and cell-based studies demonstrate that RSTS-associated AUTS2 mutations in the HX repeat domain, but not a mutation outside this region, disrupt AUTS2-P300 interaction and attenuate AUTS2-mediated active transcription. Of note, a recent study reports a patient with a syndromic neurodevelopmental disorder harboring a different mutation (532-541 aa deletion) in the AUTS2 HX repeat domain(Martinez-Delgado et al., 2020), further pointing to the critical role of AUTS2 in normal brain functioning. Moreover, mutations within exon 9 outside the HX repeat such as the PY motif, as well as a mutation at residue 495, result in individuals that display severe behavioral phenotypes such as epilepsy, in lieu of RSTS (Table S1); further stressing the role of AUTS2 in normal brain function. Consistent with the notion that ncPRC1.3 is important in the brain, mutations in FBRSL1(Ufartes et al., 2020), which binds competitively with AUTS2 to ncPRC1.3, are also associated with a neurodevelopmental syndrome. However, it is not yet known whether FBRSL1-ncPRC1.3 acts in a repressive or activating manner.

Taken together, our findings support a model in which NRF1 is a key factor that directs AUTS2-ncPRC1.3 binding to a subset of neuronal differentiation-related genes that are thereby subjected to activation by AUTS2 interaction with P300 through the AUTS2 HX repeat domain (Figure S9). As shown here, ablation of NRF1 or AUTS2 leads to defective progenitor to motor neuron differentiation *in vitro*. Deletion of NRF1 in the mouse brain and mutations in the AUTS2 HX repeat domain in humans show CC malformation, a common feature of RSTS patients which is largely associated with specific pathogenic variants in the *EP300/CREBBP* genes (Figure S9). The precise mechanisms by which NRF1 and AUTS2-ncPRC1.3 coordinately regulate gene expression, neuronal differentiation and thus mouse and human brain development and function in post-natal individuals require further investigation. Alternate approaches need to be explored as deletion of either NRF1 or AUTS2 (both long and short isoforms) apparently lead to early embryonic lethality. The generation of mouse models carrying mutations in the AUTS2 HX repeat domain might expedite future studies should they recapitulate the RSTS phenotype or other neurological diseases.

---

**Figure S1.**
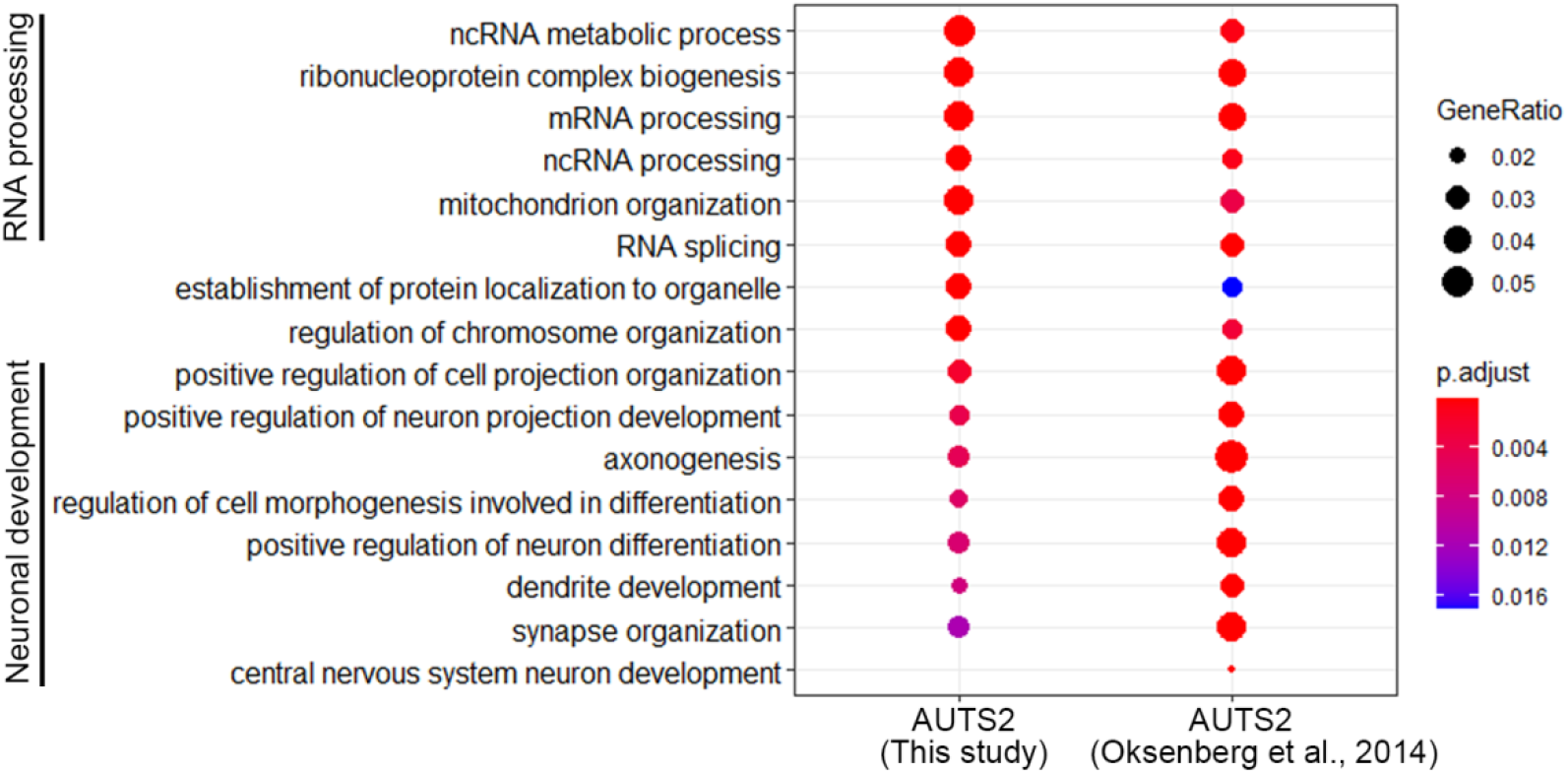
GO analysis of AUTS2-bound regions in mouse brain from our study and a study from(Oksenberg et al., 2014), related to Figure 1.

**Figure S2.**
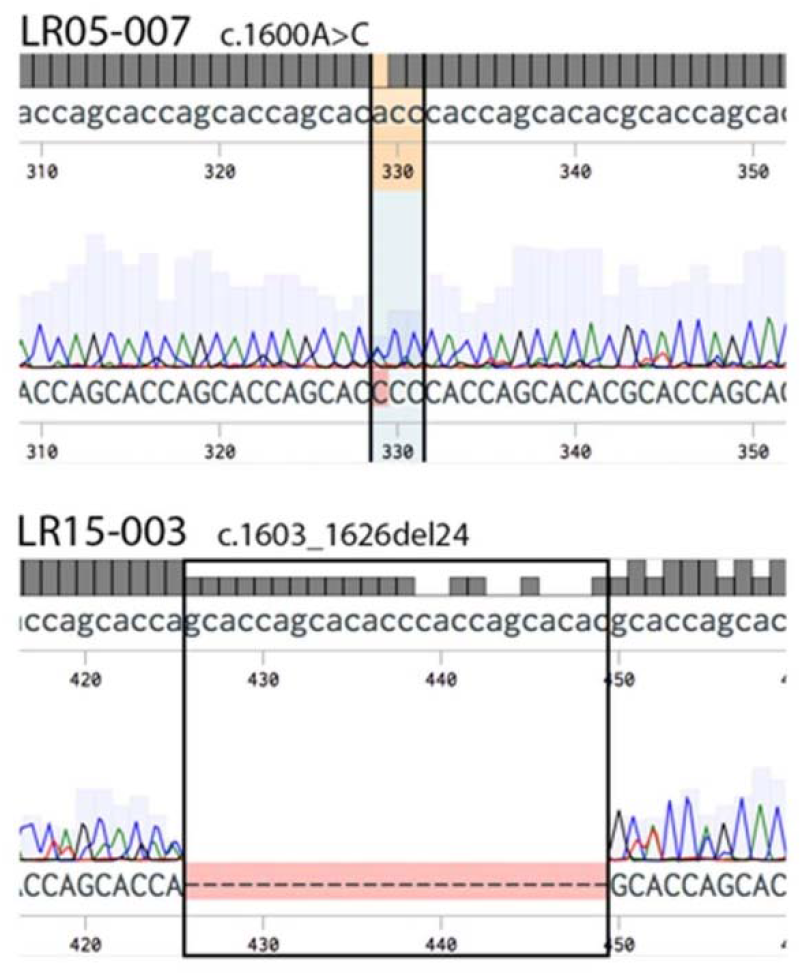
Chromatograms showing Sanger sequencing confirmation of *AUTS2* mutations in cDNA amplified from patient fibroblasts that were initially detected by exome sequencing of genomic DNA, related to Figure 2.

**Figure S3.**
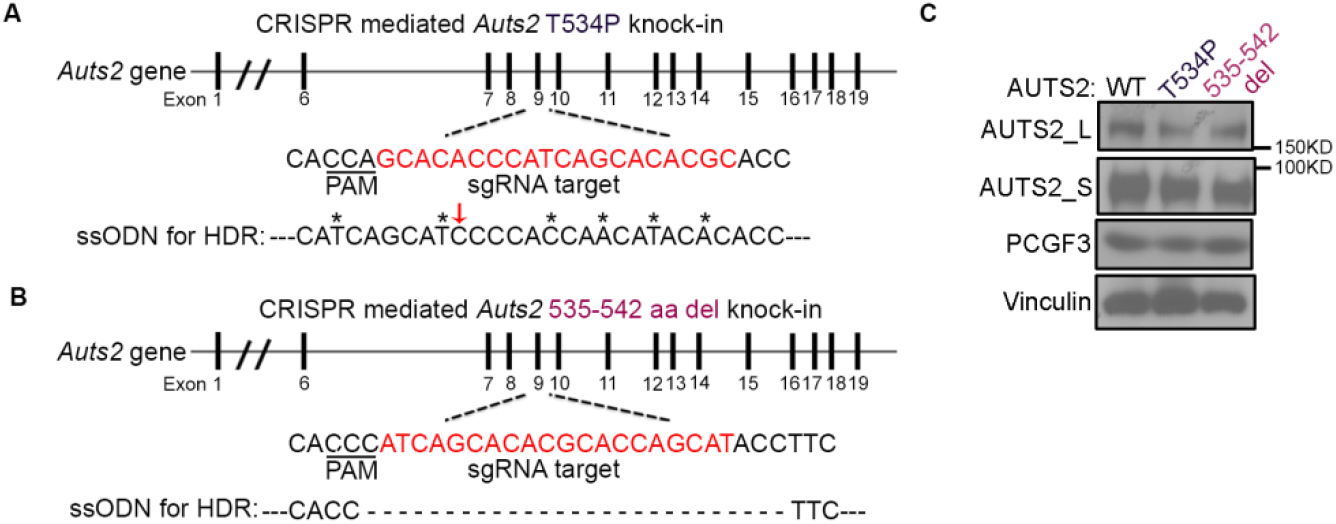
CRISPR-Cas9-mediated knock-in of mutations in AUTS2 HX repeat domain in mESC, related to Figure 3. (A) The design of sgRNA and single stranded oligodeoxynucleotide (ssODN) donor used for *Auts2* T534P knock-in with CRISPR-Cas9. Red arrow indicates T534P mutation and asterisk*-denotes the nucleotide substitution to avoid CRISPR targeting. (B) The design of sgRNA and single stranded oligodeoxynucleotide (ssODN) donor used for *Auts2* 535-542 aa deletion with CRISPR-Cas9. The 24 bps for coding AUTS2 535-542 aa were deleted in ssODN. (C) Western blot showing the expression of AUTS2 and PCGF3 in motor neuron differentiated from WT and Auts2 HX mutant (T534P and 535-542 aa deletion respectively) mESC as indicated.

**Figure S4.**
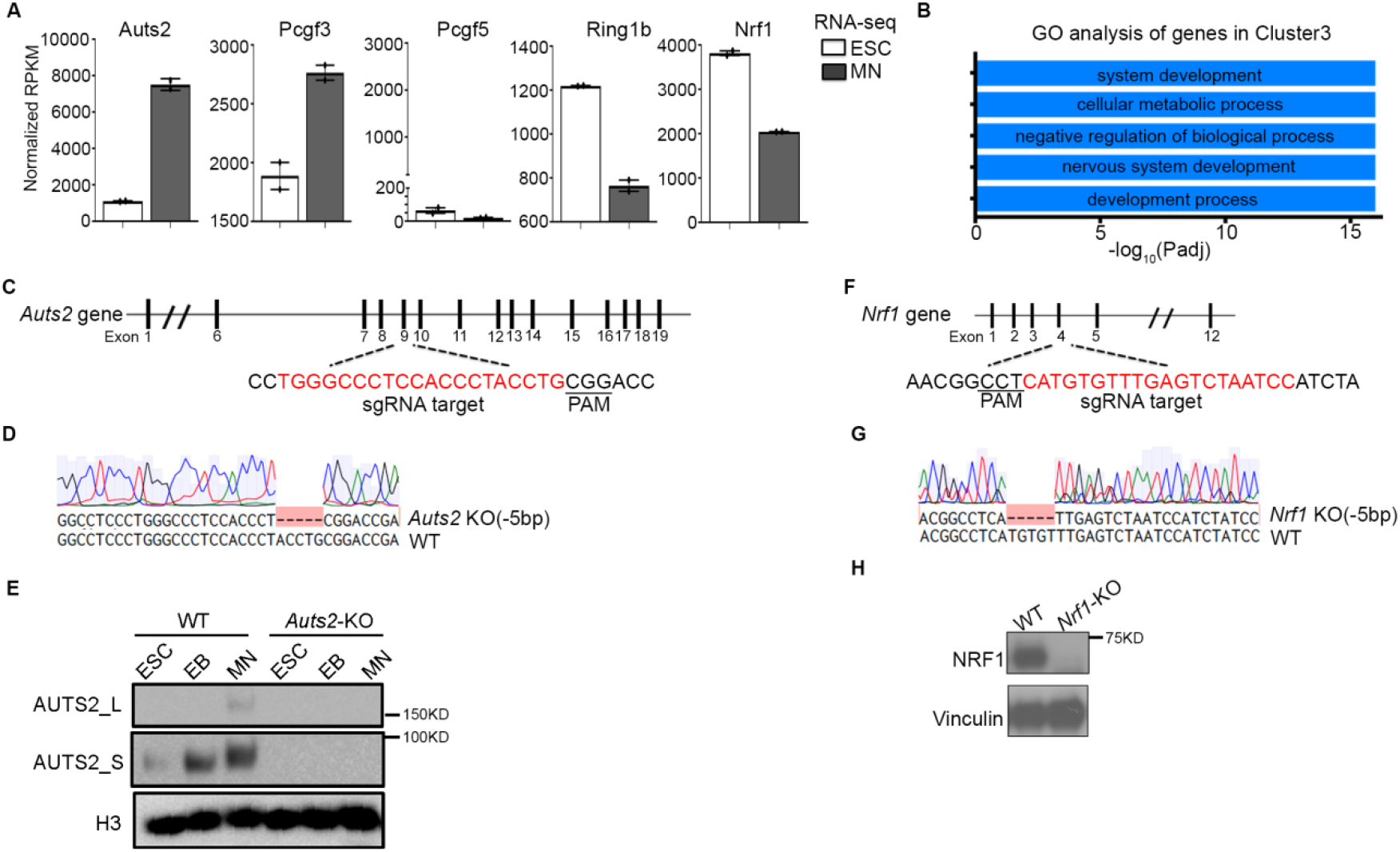
Design and validation of *Auts2* and *Nrf1* knockout, related to Figure 5. (A) Bar graphs showing the value of RPKM for Auts2, Pcgf3, Pcgf5, Ring1b and Nrf1 revealed by RNA-Seq performed on ESC and MN. (B) GO analysis of genes associated with cluster 3 regions (as indicated in Figure 3C) from Ring1B ChIP-Seq in MN. (C-E) Design (C), sanger sequencing (D) and western blot analysis (E) for CRISPR-Cas9 mediated *Auts2* knockout. (F-H) Design (F), sanger sequencing (G) and western blot analysis (H) for CRISPR-Cas9 mediated *Nrf1* knockout.

**Figure S5.**
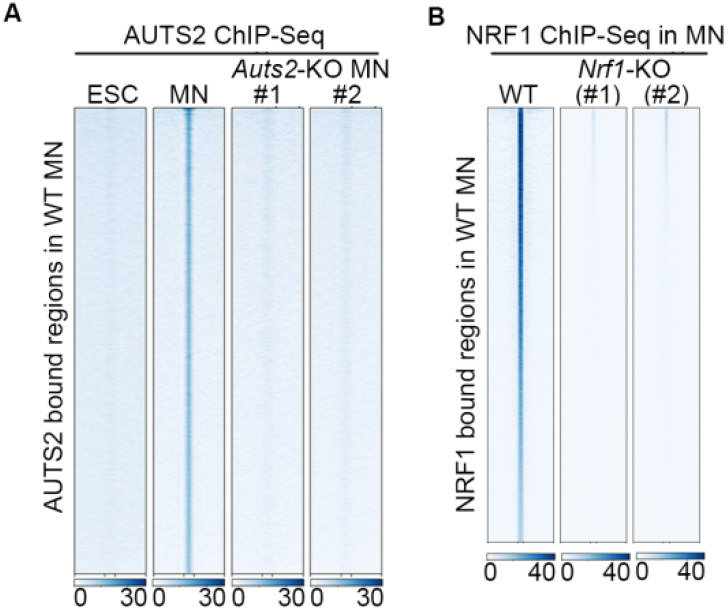
Validation of AUTS2 and NRF1 ChIP-seq specificity under *Auts2*-KO and *Nrf1*-KO, related to Figure 5. (A) Heatmap showing AUTS2 ChIP-seq signals from ESC and either WT or *Auts2*-KO MN centered on AUTS2-bound regions identified in WT MN (±5 kb). #1 and #2 represent 2 *Auts2*-KO clones. (B) Heatmap showing NRF1 ChIP-seq signals from WT and *Nrf1*-KO ESC centered on NRF1-bound regions identified in WT ESC (±5 kb). #1 and #2 represent 2 *Nrf1*-KO clones.

**Figure S6.**
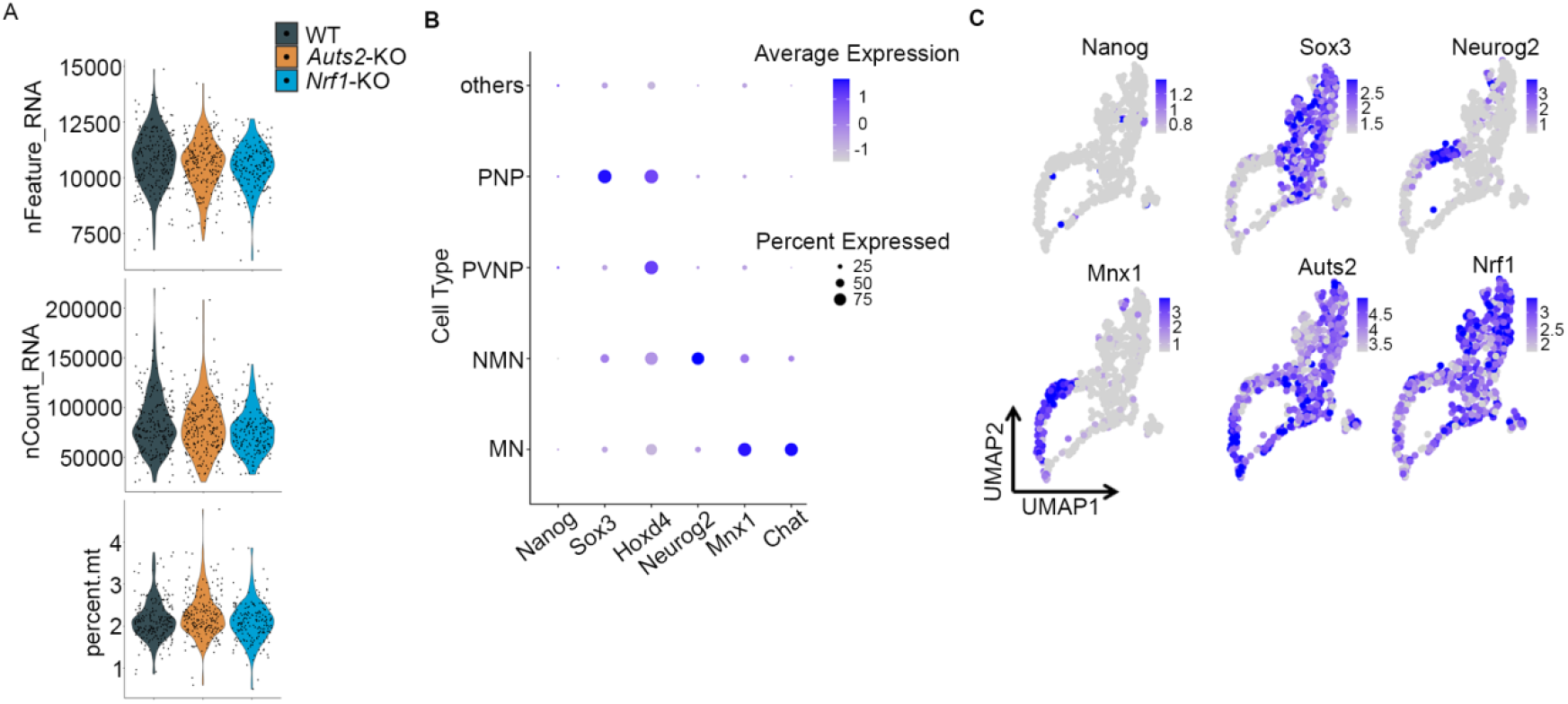
Quality control for scRNA-seq, related to Figure 6. (A) Violin plots summarizing the number of genes, molecules detected and percentage of reads mapping to mitochondrial genome per cell by scRNA-seq in WT, *Auts2*-KO and *Nrf1*-KO samples. (B) Dot plot shows the expression levels and frequencies of cell-type-specific marker genes in each cluster. The size and color of the dot represent the percentage of cells expressing the marker gene and the relative expression level (Z-score) within a cell cluster, respectively. (C) UMAP plots are colored by the expression level of the genes indicated. *Nanog*: stem cell marker, *Sox3*: posterior neural progenitor marker, *Neurog2*: newborn motor neuron marker, and *Mnx1*: motor neuron marker.

**Figure S7.**
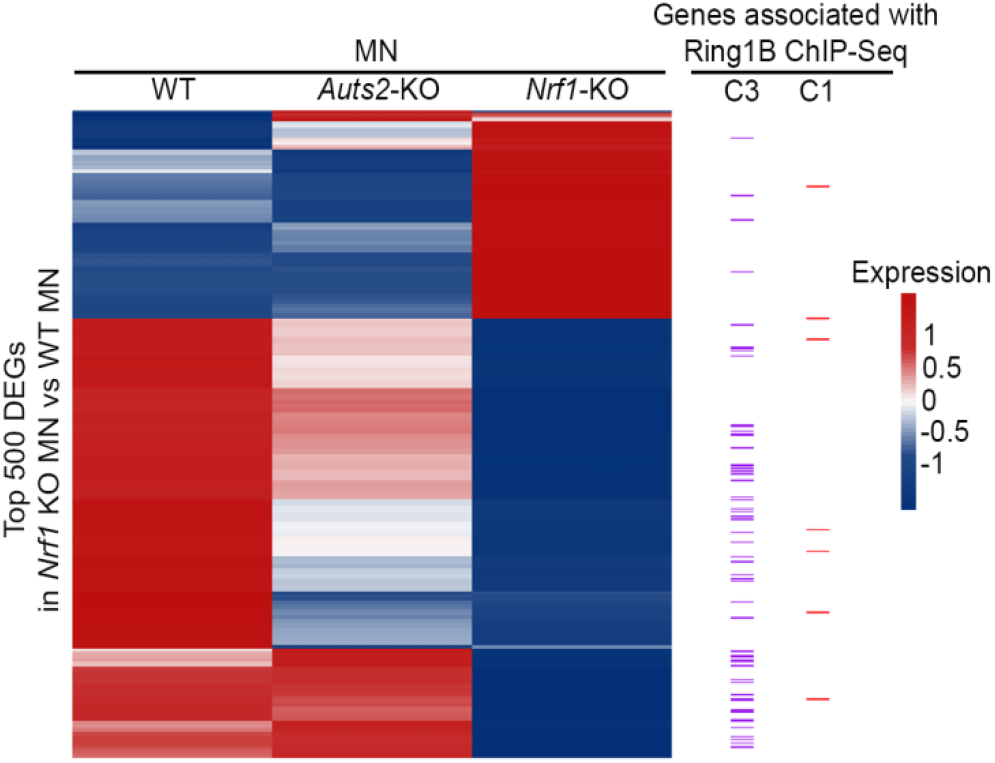
Comparing the differentially expressed genes in *Nrf1*-KO MN by scRNA-Seq with PRC1 associated active and repressed genes revealed by Ring1B ChIP-Seq, related to Figure 6. Top 500 differentially expressed genes (DEGs) were identified by comparing *Nrf1*-KO MN versus WT MN. Expression of these DEGs across WT MN, *Auts2*-KO MN and *Nrf1*-KO MN is shown by heatmap. Color scale represents the averaged and scaled expression values from each cell population. Each row represents a gene and whether the gene is located in cluster 1 or cluster 3 regions from Ring1B ChIP-Seq (see Figure 5C for more details) is marked on the right.

**Figure S8.**
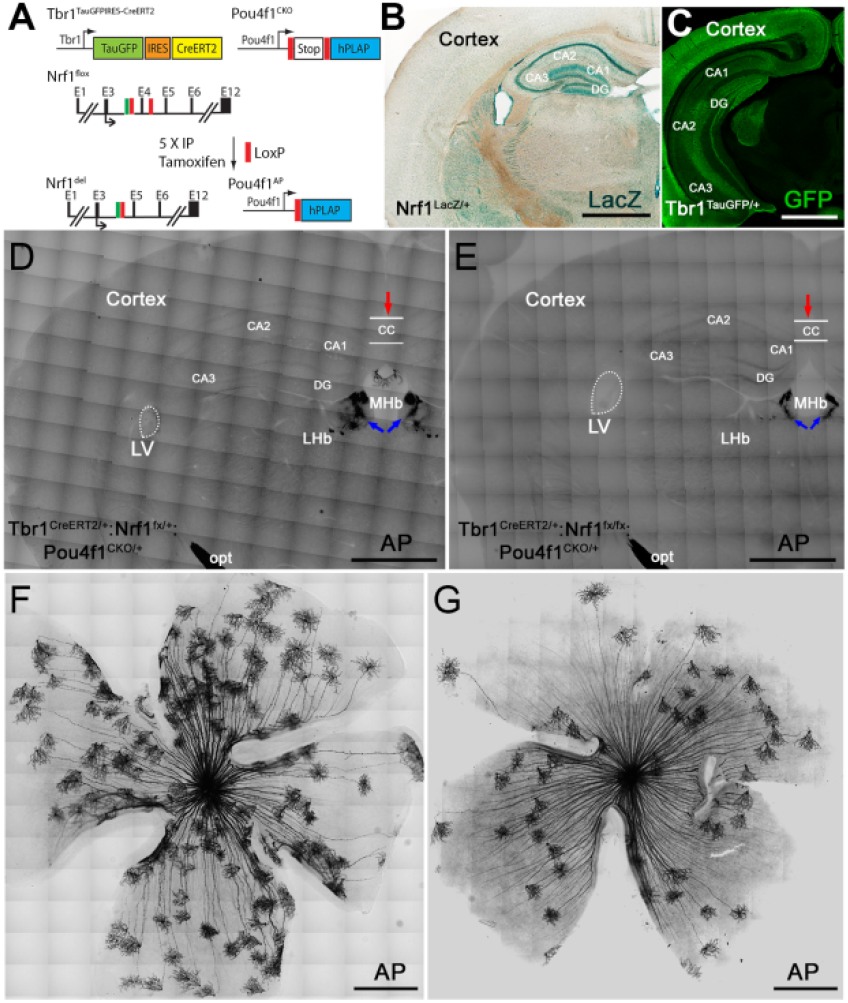
*Tbr1^CreERT2^*-mediated *Nrf1*-deletion in CNS leading to neuronal loss. (A) Schematic illustration showing the strategy for *Tbr1^CreERT2^*-mediated conditional KO of exon 4 of the *Nrf1* gene. Expression of TauGFP and Cre^ERT2^ is driven by the Tbr1 promoter. Double loxP sites were inserted into the third and fourth introns of the *Nrf1* gene. Double loxP sites spanning a translation stop codon and followed by the coding region of human placental alkaline phosphatase (AP) were also inserted into the 5’ UTR region of the *Pou4f1* gene. After Cre^ERT2^ activation by tamoxifen, *Nrf1* exon 4 will be deleted and hPLAP will be expressed and used for AP staining. (B) NRF1 expression pattern in adult brain revealed by LacZ expression in the *Nrf1^LacZ/+^* mouse. (C) Tbr1 expression pattern as revealed by immunostaining using anti-GFP antibody on *Tbr1^TauGFP/+^* brain section. (D-E) Representative images of brain sections (at Bregma −1.7 mm) from control (D, *Tbr1^CreERT2/+^:Nrf1^fx/+^:Pou4f1^CKO/+^*) and mutant (E,*Tbr1^CreERT2/+^:Nrf1^fx/fx^:Pou4f1^CKO/+^*) mice. Brain sections were collected three months after intraparietal injection of tamoxifen. A significant reduction in the size of the hippocampus (CA1, CA2, CA3, DG), the width of the corpus callosum (CC, red arrow) and the number of alkaline phosphatase (AP)+ fibers passing through the medium and lateral habenula (MHb, blue arrows), and a significant enlarged lateral ventricle (dotted circle) were observed in the mutant brain section, indicative of neuronal loss upon *Nrf1* knockout. CA1-3: hippocampus sub-regions; DG: dentate gyrus. (F-G) Representative AP-stained images showing *Tbr1*-expressing RGC in retinas from control (F, *Tbr1^CreERT2/+^: Nrf1^fx/+^: Pou4f1^CKO/+^*) and mutant (G, *Tbr1^CreERT2/+^: Nrf1^fx/fx^: Pou4f1^CKO/+^*) mice. Retinas were collected three months after intraparietal injection of tamoxifen. The number of AP+ RGC in the mutant retinas was reduced by approximately 45% compared to control (N = 4; control: 112.25 ± 8.30, mutant: 63.5 ± 11.7; P = 0.0005), further confirming neuronal loss in the central nervous system upon *Nrf1* knockout. Scale bars: 1 mm in B-E, 500 μm in F and G.

**Figure S9.**
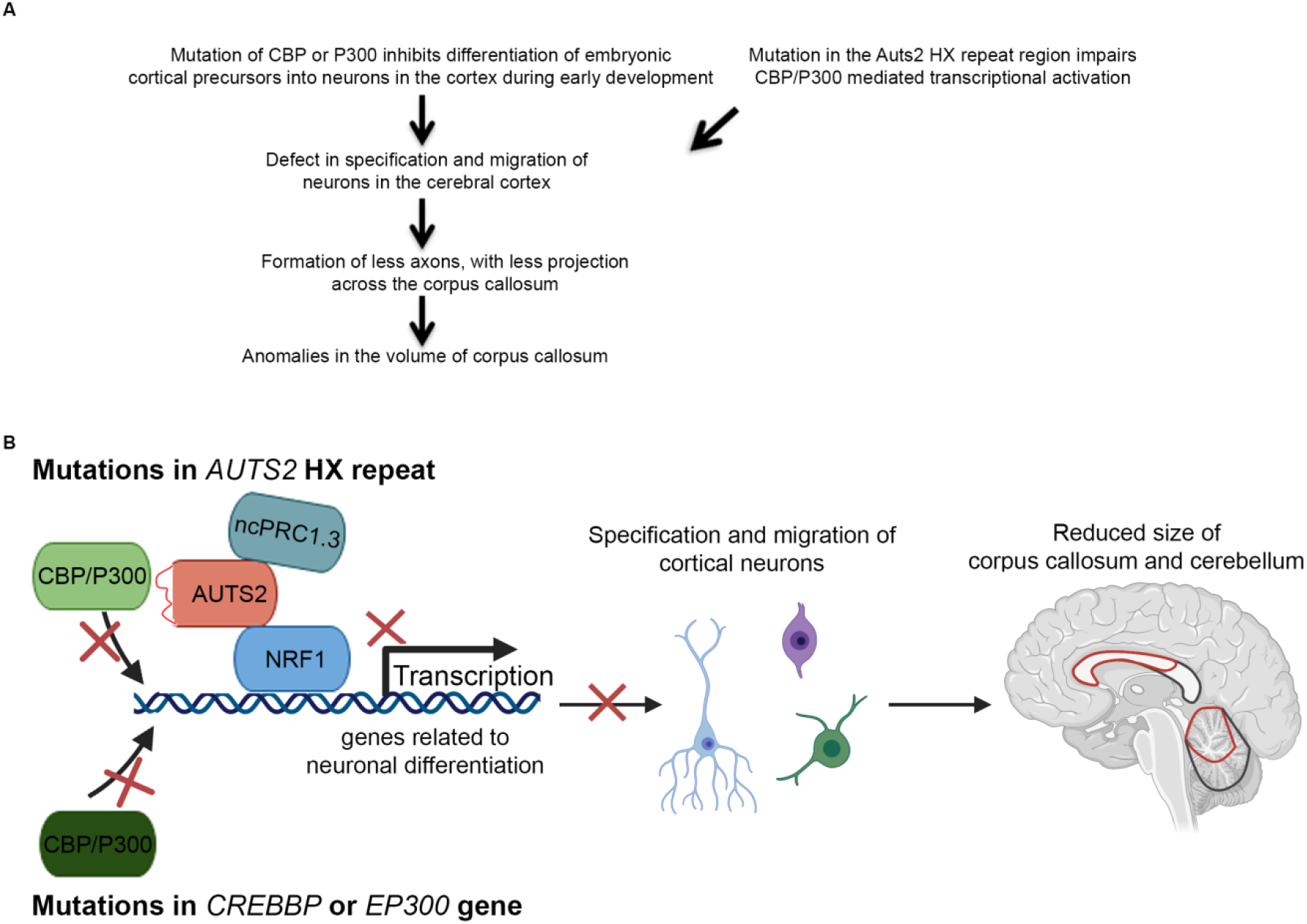
Hypothetical model for associating mutations in *AUTS2* HX repeat with Rubinstein-Taybi syndrome.

## STAR*METHODS

### KEY RESOURCES TABLE

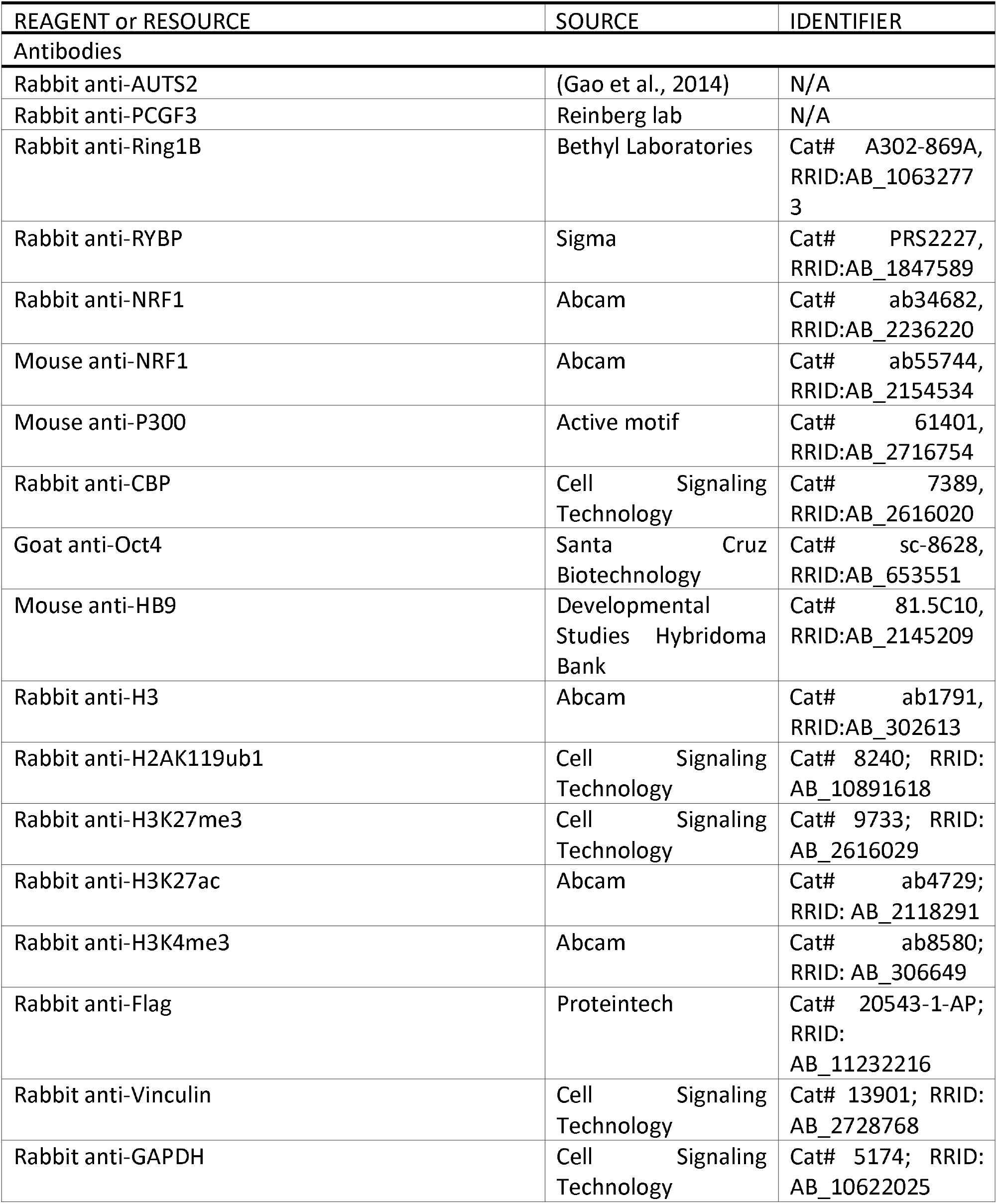
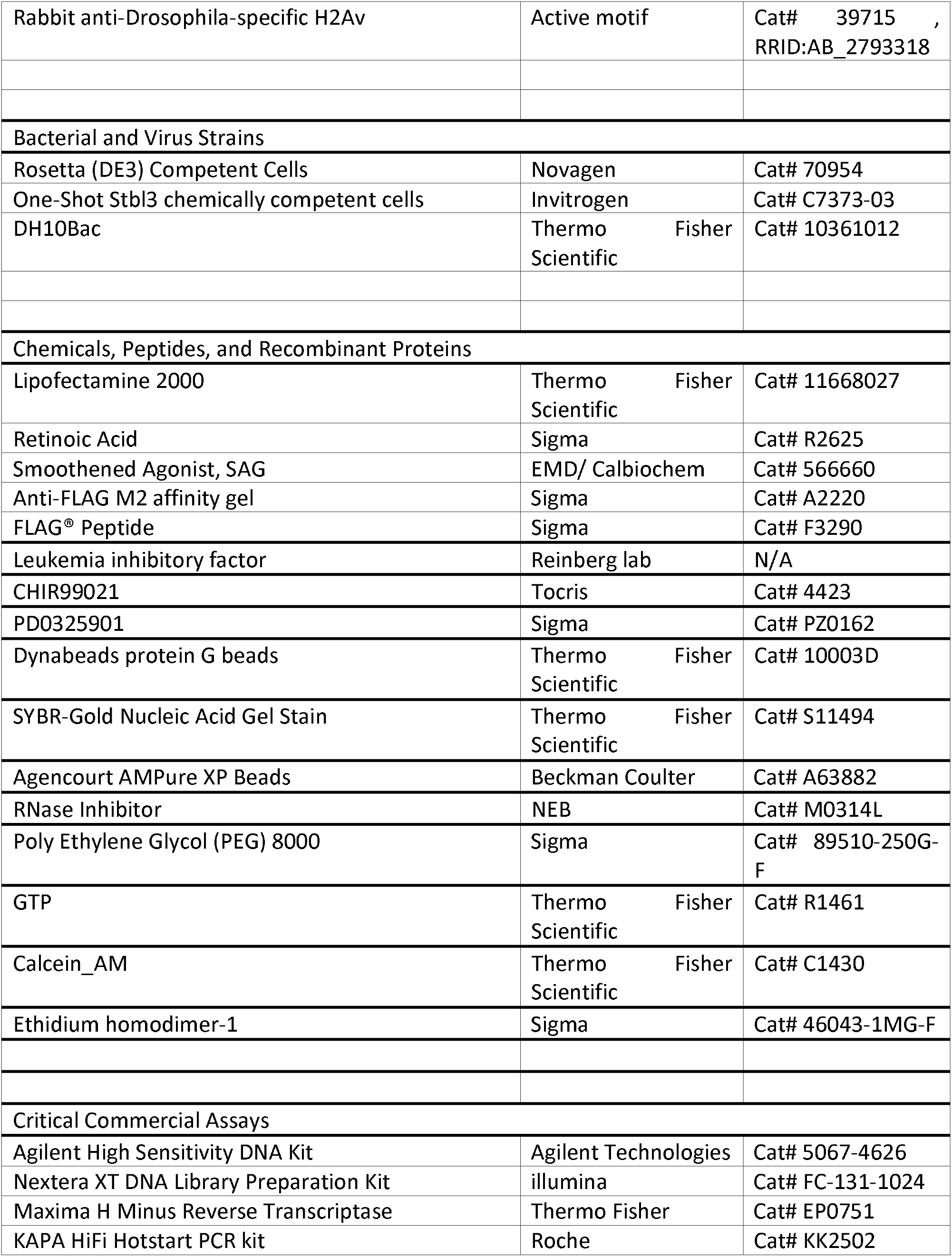
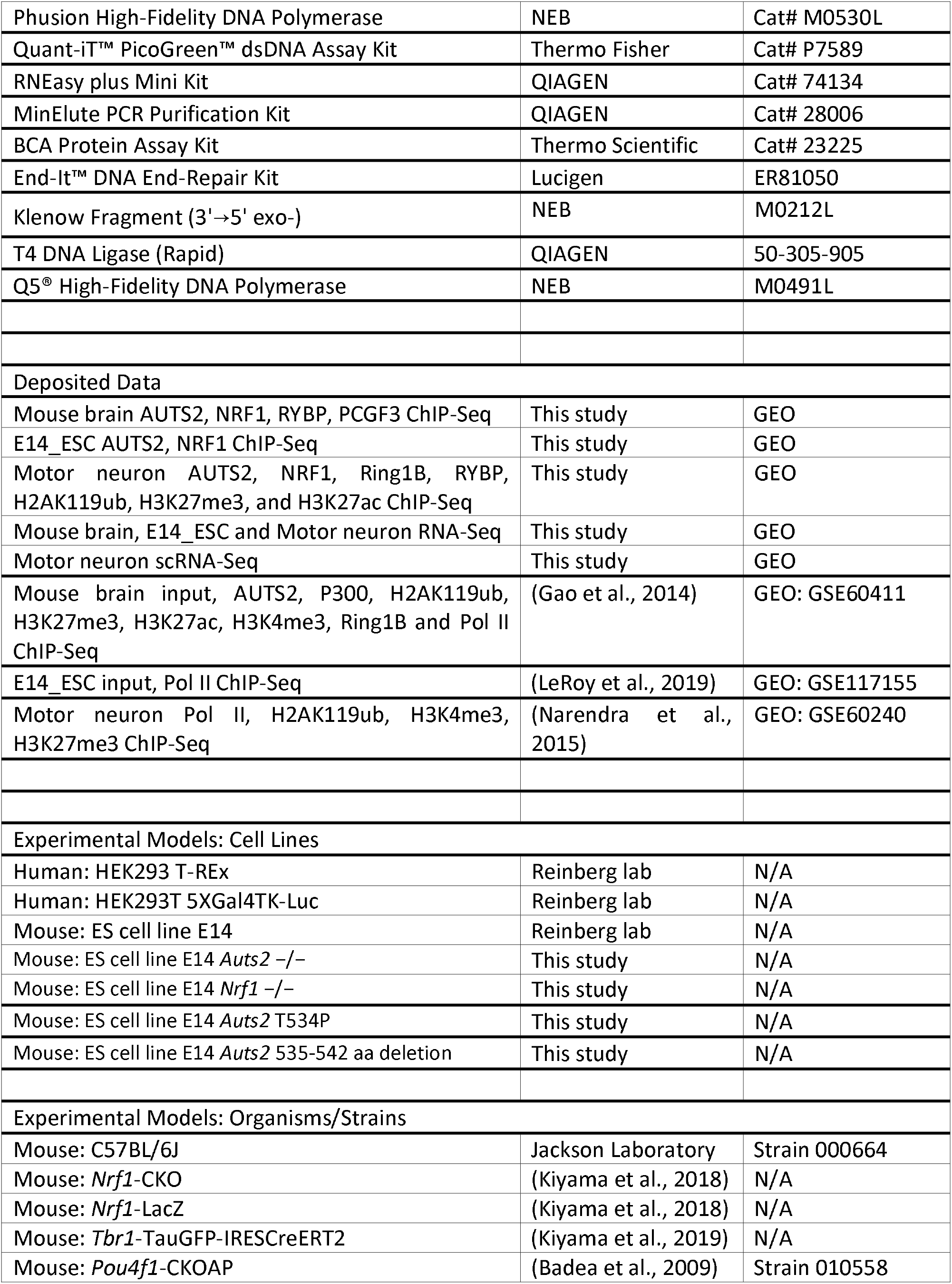
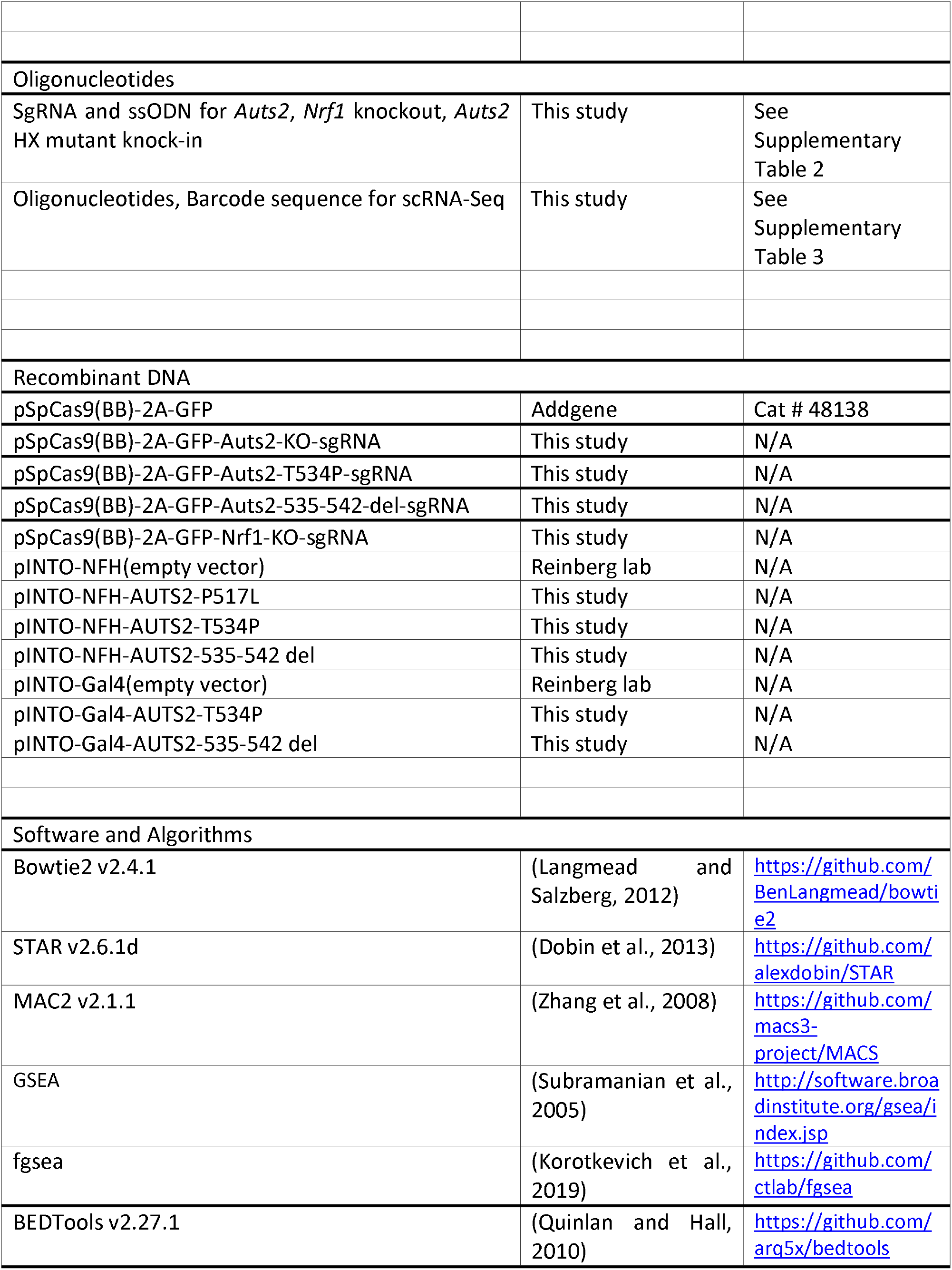
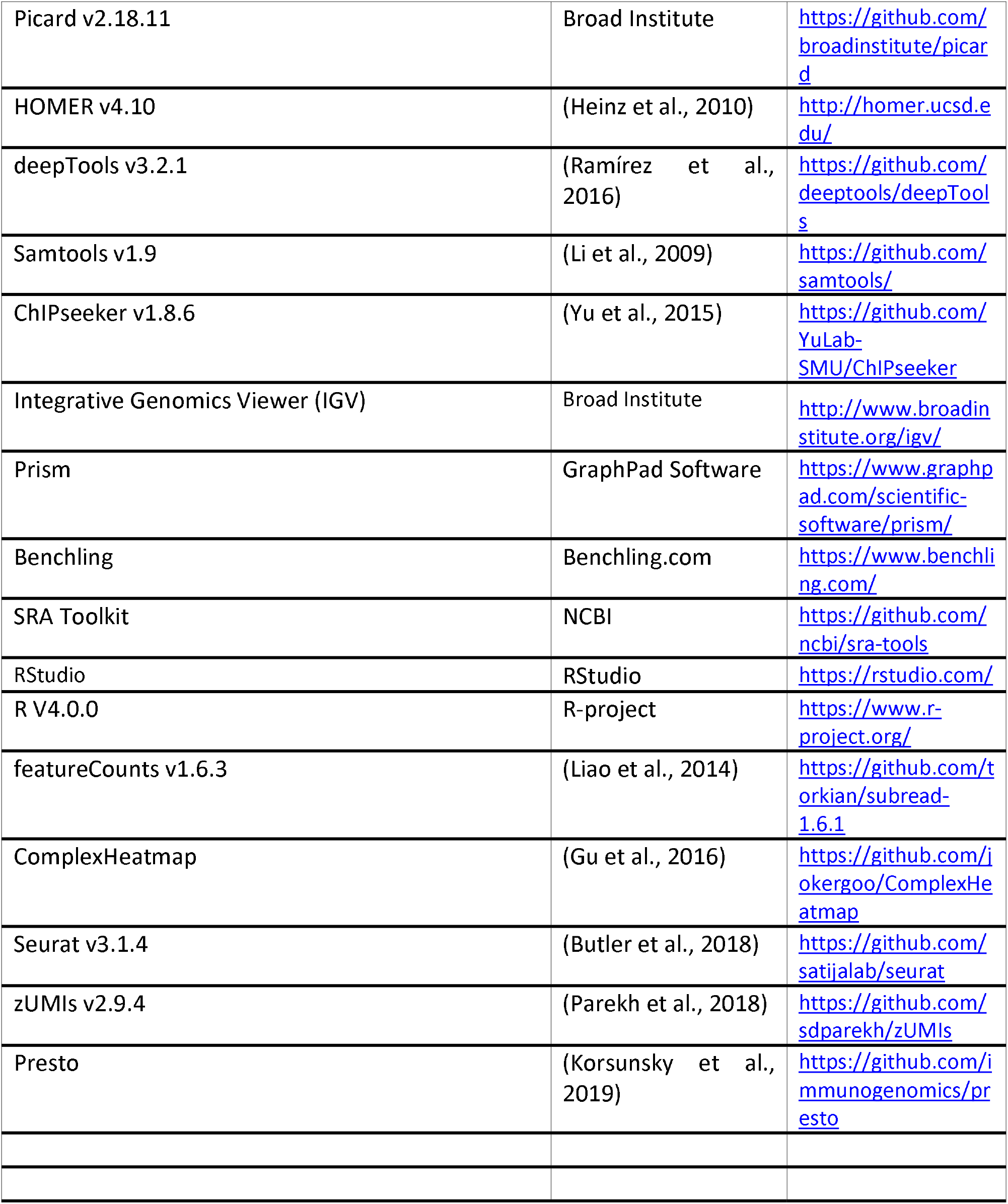

#### Lead Contact and Materials Availability

Please direct inquiry to the Lead Contact, Danny Reinberg. All unique/stable reagents generated in this study are available from the Lead Contact with a completed Materials Transfer Agreement.

#### Experimental Model and Subject Details

##### Animals

All mice were housed with a 12 hour light-dark cycle. Mixed cohorts of female and male mice were used for all experiments to minimize gender effects. All animal procedures followed the US Public Health Service Policy on Humane Care and Use of Laboratory Animals and were approved by the Animal Welfare Committee at New York University and the University of Texas McGovern Medical School at Houston.

##### Cell lines and culture condition

All ESC lines (E14 and derivatives) were grown in DMEM supplemented with 15% FBS, L-glutamine, penicillin/streptomycin, sodium pyruvate, non-essential amino acids, 0.1 mM β-mercaptoethanol, LIF, and 2i inhibitors, which include 1 μM MEK1/2 inhibitor (PD0325901) and 3 μM GSK3 inhibitor (CHIR99021) on 0.1% gelatin coated plates.

HEK293 T-Rex and HEK293T 5XGal4TK-Luc cells were cultured in standard DMEM supplemented with 10% FBS, 100 U/mL penicillin-streptomycin.

WT and mutant NFH-AUTS2 inducible cell lines were obtained by transfecting each pINTO-NFH plasmid into 293 T-Rex cells, and WT and mutant Gal4-AUTS2 inducible cell lines were obtained by transfecting each pINTO-Gal4 plasmid into HEK293T 5XGal4TK-Luc cells. Transfected cells were seeded at limiting dilutions, and isolated clones were screened by western blot.

##### Clinical Cohort

The initial proband (LR05-007) was identified through trio-based exome sequencing in a cohort of 100 individuals with cerebellar malformations(Aldinger et al., 2019). We recruited 6 additional individuals with *de novo* pathogenic or likely pathogenic variants in *AUTS2* by sharing data through GeneMatcher(Sobreira et al., 2015) or through collaboration with colleagues. The 7 individuals in this cohort (2 females, 5 males) were 1 to 15.5 years of age at the time of their most recent evaluation. We obtained clinical data for all patients, including features tabulated in a reported AUTS2 clinical severity score(Beunders et al., 2013, 2015, 2016). Written informed consent was obtained from all participants through protocols approved by Institutional Review Boards at the local institution or at Seattle Children’s Hospital.

## METHOD DETAILS

### Immunoprecipitation and Proteomics

Cell pellets were prepared from cell culture plates or mouse brain. Nuclei were extracted using HMSD buffer (20 mM HEPES, pH 7.5 at 4°C, 5 mM MgCl_2_, 250 mM sucrose, 25 mM NaCl, 1 mM DTT) supplemented with protease inhibitors (0.2 mM PMSF, 1 μg/mL Pepstatin A, 1 μg/mL Leupeptin, and 1 μg/mL Aprotinin) and phosphatase inhibitors (10 mM NaF and 1 mM Na_3_VO_4_), and incubated on ice for 5 min. Lysates were pelleted at 3,000 rpm for 5 min at 4°C and nuclei pellets were washed one more time with HMSD buffer. The resulting nuclei pellets were resuspended in BC420 high salt buffer (20 mM Tris-HCl at pH 7.9, 1.5 mM MgCl_2_, 0.42 M NaCl, 0.5 mM DTT, 0.2 mM EDTA) supplemented with protease inhibitors and phosphatase inhibitors for lysing at 4°C for 1 hr with occasional pipetting. Lysates were then pelleted at 20,000 x g for 15 min at 4°C. Finally, supernatants were collected and subjected to dialysis in Buffer D (20 mM HEPES, 150 mM NaCl, 1.5 mM MgCl_2_, 0.2 mM EDTA, and 5% glycerol) overnight at 4°C. Prior to any subsequent applications, nuclear extracts were centrifuged again at 20,000 g for 10 min at 4°C to remove any precipitate. Supernatants were collected, and protein concentrations were quantified via bicinchonic acid (BCA) assay. For immunoprecipitation, 1-2 mg of nuclear extract was incubated with 1~3 μg of antibody. After incubation at 4°C for 2 hr, 30 ul of protein G beads were added for incubation at 4°C overnight. Beads were washed three times with Buffer D (20 mM HEPES, 150 mM NaCl, 1.5 mM MgCl_2_, 0.2 mM EDTA, and 5% glycerol), and eluted with 0.2 M glycine (pH 2.6) or 1× SDS loading buffer. Proteins from immunoprecipitation were separated by SDS–PAGE, using 4%–12% NuPAGE Novex Bis–Tris gels and then stained with Coomassie Blue. Bands were excised from gels and digested with trypsin, followed by standard LC-MS/MS procedure.

### Whole cell extract and western blotting

Cells were harvested and lysed with CHAPS-Urea buffer (50 mM Tris-HCl, pH 7.9, 8M Urea, and 1% CHAPS) containing protease inhibitors and phosphatase inhibitors as mentioned above. The cell suspension was briefly sonicated (40% amplitude, 5 strokes) and centrifuged at 20,000 x g at 4°C for 20 min. The supernatant was collected and protein concentrations were quantified via bicinchonic acid (BCA) assay. Proteins were separated using a 6%–12% SDS PAGE gel, and transferred onto a PVDF membrane. Membranes were blocked with 5% milk in PBST at RT for 1 hr and incubated with primary antibody overnight at 4°C. Membranes were washed 3 times with PBST and then incubated with HRP-conjugated secondary antibodies for 1 hr at RT, followed by exposure.

### CRISPR-mediated genome editing

To generate stable *Auts2* and *Nrf1* KO cell lines, sgRNAs were designed using CRISPR design tool in https://benchling.com. sgRNAs in Table S1 were cloned in pSpCas9(BB)-2A-GFP (PX458, a gift from Feng Zhang, Addgene plasmid #48138) and transfected into mESCs, using Lipofectamine 2000 (Life Technologies). GFP-positive cells were sorted 48 hr after transfection and 20,000 cells were plated on a 15 cm dish. Single mESC was allowed to grow to a colony for ~5 days and then was picked, trypsinized in Accutase for 5 min, and split into two individual wells of two 96-well plates for genotyping and culture, respectively. Genomic DNA was extracted using lysis buffer (50 mM Tris-HCl, pH 8, 2 mM NaCl, 10 mM EDTA, 0.1% SDS) supplemented with protease K, and genotyping PCRs were performed using primers (Table S2) surrounding the target site. The resulting PCR products were sent for sequencing to determine the presence of a deletion or a mutation event. Clones were further confirmed by western blot.

For endogenously knock-in the mutations in AUTS2 HX repeat in mESC, a ssODN donor used for homology directed repair and CRISPR/Cas9 plasmid (px458) with designed sgRNA (see Table S2 also) were co-transfected into mESCs, using Lipofectamine 2000. The following FACS, colony picking and characterization are the same as generating KO lines described above.

### Motor Neuron Differentiation

E14 mouse embryonic stem cells (mESCs) were cultured in standard medium supplemented with LIF, and 2i conditions as described above. For motor neuron (MN) differentiation, the previously described protocol was applied(Narendra et al., 2015). Briefly, about 4 million mESCs were plated in a 500 cm^2^ square dish and differentiated into embryoid bodies in AK medium (250 ml advanced DMEM/F12, 250 ml neurobasal medium, 75 ml knockout serum, L-glutamine, penicillin/streptomycin, 0.1 mM β-mercaptoethanol) for 2 days. Embryoid bodies were then diluted by 1:4 and further patterning was induced by freshly adding 1 μM all-trans-retinoic acid (RA) and 0.5 μM smoothened agonist (SAG) for an additional 4 days. Fresh medium was added after 2 days to support motor neuron survival.

### Luciferase reporter assay

HEK293T 5XGal4 TK-Luc cells stably transfected with pINTO-GAL4 vector control or with inserts of interest were treated with 100 ng/ml doxycycline. Cells were lysed by adding 250 ul of ice-cold lysis buffer (0.2% Triton X-100, 100 mM potassium phosphate, pH 7.8, and 1 mM DTT) and shaking for 10 min at 4°C. The cell lysate was centrifuged at 20,000g for 10 min and the protein concentration of the resulting supernatant was determined by bicinchonic acid (BCA) assay. 30 ug of the supernatant was assayed for luciferase activity using luciferase assay substrate (Promega).

### X-gal staining

Mouse brain was fixed by perfusion with 10% neutral buffered formalin. Extracted brain was embedded in OCT compound then sectioned into 50 um thickness. Sections were dried at RT for 3 hr and then washed with wash buffer (0.1 M sodium phosphate containing 2 mM MgCl_2_, 0.01% deoxycholate, and 0.02% Nonidet P-40, pH 7.3). LacZ color reaction was performed in wash buffer containing 5 mM potassium ferrocyanide, 5 mM potassium ferricyanide, and 1 mg/ml X-gal at 37°C overnight. Color reaction was terminated by incubation in 10% formalin for 10 min. Post-fixed sections were washed, dehydrated, and mounted with Cytoseal 60 (Thermo Fisher Scientific). Images were collected with a Canon EOS 10 digital camera (Melville, NY) mounted on an Olympus IX71 microscope.

### Immuno-histochemical analysis

Mouse brain was fixed by perfusion with 10% neutral buffered formalin. Extracted brain was embedded in OCT compound, and then sectioned into 100 um thickness. Sections were incubated with anti-GFP (1:1000, Invitrogen) antibody. Alexa-488 conjugated secondary antibody was used in 1:800 dilution (Jackson Immuno-reserach).

### Alkaline phosphatase (AP) staining

*Tbr1^CreERT2/+^:Nrf1^fx/+^:Pou4f1^CKO/+^* and *Tbr1^CreERT2/+^:Nrf1^fx/fx^:Pou4f1^CKO/+^* mice were used for AP staining 3 months after tamoxifen injection. Brains were fixed by perfusion with 10% neutral buffered formalin. Fixed brain was embedded in OCT compound and sectioned into 100 um thickness. Retinal flat-mounts were fixed in 10% neutral buffered formalin for 10 min at RT. Brain sections and retinas were incubated in heated PBS for 30 min in a 65 C water bath to inactivate endogenous AP activity. AP color reaction was performed in 0.1 M Tris, pH 9.5, 0.1 M NaCl, 50 mM MgCl_2_, 0.34 g/ml nitroblue tetrazolium and 0.175 g/ml 5-bromo-4-chrolo-3-indolyl-phosphate for overnight at RT. Stained tissues were washed three times in PBS, post-fixed with 10% neutral buffered formalin, dehydrated with a series of ethanol, then cleared with 2:1 benzyl benzoate/benzyl alcohol. Tiled images were collected using a Zeiss Axio Imager 2 microscope (Carl Zeiss).

### ChIP-seq library preparation

For cross-linking, ESCs were fixed in 1% formaldehyde for 10 min at RT directly on plates and quenched with 125 mM glycine for 5 min at RT. For cross-linking of MN, ESC-derived motor neuron cultured for 6 days were dissociated with 0.05% trypsin, neutralized, fixed in 1% formaldehyde for 15 min at RT and then quenched with 125 mM glycine for 5 min at RT. For cross-linking of mouse brain, mouse whole brains were quickly dissected at postnatal day one and homogenized with a glass douncing homogenizer using first a loose, then a tight pestle. The cell homogenate was fixed with a final concentration of 1% formaldehyde for 10 min at RT and the reaction was quenched with 0.125 M glycine for 5 min at RT.

Cell pellets were washed twice in PBS and nuclei were isolated using buffers in the following order: LB1 (50 mM HEPES, pH 7.5 at 4°C, 140 mM NaCl, 1 mM EDTA, 10% Glycerol, 0.5% NP40, 0.25% Triton X; 10 min at 4°C), LB2 (10 mM Tris, pH 8 at 4°C, 200 mM NaCl, 1 mM EDTA, 0.5 mM EGTA; 10 min at 4°C), and LB3 (10 mM Tris, pH 7.5 at 4°C, 1 mM EDTA, 0.5 mM EGTA, and 0.5% N-Lauroylsarcosine sodium salt). Chromatin was fragmented to an average size of 250 bp in LB3 buffer using a Diagenode Bioruptor. 200 μg sonicated chromatin, 4 ug antibody and 20 ul Dynabeads were used in each ChIP reaction supplemented with 0.5x volumn of incubation buffer (3% Triton X, 0.3% Na Deoxycholate, 15 mM EDTA). 1 μg of Drosophila chromatin and 0.2 μg of anti-Drosophila H2A.X antibody were added in each ChIP reaction as spike-in references, except 3 μg of Drosophila chromatin was used for H3K27ac and H2AK119ub ChIP. After 5 consecutive washes with RIPA buffer (50 mM HEPES, pH 7.5 at 4°C, 0.7% Na Deoxycholate, 1 mM EDTA, 1% NP40, 500 mM LiCl) and one wash with TE+50 mM NaCl, the beads-bound DNA was eluted in freshly prepared elution buffer (50 mM Tris, pH 8, 10 mM EDTA, 1% SDS) at 65°C for 20 min. Eluted DNA was de-crosslinked at 65°C overnight, followed by protease K and RNase A treatment.

For Library preparation, IP’ed DNA (~1-30 ng) was end-repaired using End-It Repair Kit, tailed with deoxyadenine using Klenow exo-, and ligated to custom adapters with T4 Rapid DNA Ligase (Enzymatics). Fragments of 200-600 bp were size-selected using Agencourt AMPure XP beads (0.5X and 0.3X), and subjected to PCR amplification using Q5 DNA polymerase. Libraries were size-selected using Agencourt AMPure XP beads (0.75X), quantified by Qubit™ dsDNA HS Assay Kit and quality checked by High Sensitivity D1000 ScreenTape. Libraries were sequenced as 50 bp single-end reads on the Illumina HiSeq 4000 platform.

### RNA-seq library preparation

Total RNA was isolated with Tripure isolation reagent and gDNA was removed by RNeasy Plus mini kit. PolyA+ RNA was isolated from 5 ug total RNA using Dynabeads Oligo(dT)_25_, fragmented with Mg^2+^ contained in the 1^st^ strand buffer at 94°C for 15 min, and reverse transcribed using Superscript III and random hexamers to synthesize the first strand cDNA. Single strand cDNA was precipitated and dTTP was removed by phenol/chloroform/isoamyl alcohol extraction and ethanol precipitation. Second strand cDNA was synthesized with dUTP to generate strand asymmetry using DNA Pol I and E. coli ligase, and then purified by MinElute PCR Purification Kit. Double-stranded DNA was end-repaired, A-tailed, and ligated to custom barcode adapters as described above. RNA-seq libraries were sequenced as 50 bp paired-end reads on the Illumina HiSeq 4000 platform or NovaSeq 6000 platform.

### Single cell RNA-seq library preparation

Single cell RNA-seq library by Smart-seq3 technique was generated according to the published protocol(Hagemann-Jensen, 2020; Hagemann-Jensen et al., 2020), with the following modifications. ESC-derived motor neuron cultured for 6 days were dissociated with 0.05% trypsin and stained with 0.2 uM Calcein AM and 8 uM Ethidium homodimer-1 at RT for 15 min. Single viable cells were sorted using Fluorescence Activated Cell Sorting to single wells of 96-well fully-skirted Eppendorf PCR plates in 3 uL lysis buffer (5% PEG8000, 0.1% Triton X-100, 0.5 unit/ul RNAse Inhibitor, 0.5 uM OligodT_30_VN, 0.5 mM dNTP in nuclease free water). The plates were immediately covered, spun at 2000 rpm for 1 min at 4°C, and stored at −80°C until further analysis. In each plate, well A1 was left empty and 100 cells were sorted to well H1 for quality control and they were excluded for downstream analysis.

Plates were incubated at 72°C for 10⍰min for lysing the cells and denaturing the RNA. Next, 1⍰μl of reverse transcription mix (25⍰mM Tris-HCl, pH 8, 30⍰mM NaCl, 1⍰mM GTP, 2.5⍰mM MgCl_2_, 8⍰mM DTT, 0.5 unit/ul RNAse Inhibitor, 2⍰μM Smart-seq3 TSO, 2⍰ unit/ul Maxima H-minus reverse transcriptase enzyme) was added to each well for reverse transcription and template switching at 42°C for 90⍰min followed by 10 cycles at 50°C for 2⍰min and at 42°C for 2⍰min and the reaction was inactivated at 85°C for 5⍰min. cDNA pre-amplification was performed by adding 6⍰μl of PCR mix (1×Kapa HiFi HotStart buffer, 0.02⍰ unit/ul KAPA HiFi DNA polymerase, 0.5⍰mM MgCl_2_, 0.3⍰mM dNTPs, 0.5⍰μM Smartseq3 forward PCR primer and 0.1⍰μM Smartseq3 reverse PCR primer) with the following protocol: 3⍰min at 98°C for initial denaturation, 18–24 cycles of 20 s at 98°C, 30⍰s at 65°C and 6⍰min at 72°C, followed by a final elongation for 5⍰min at 72°C. PCR cycles were determined for each cell type by prior experiments and 19 cycles were used for motor neuron to obtain ~10 ng purified cDNA.

Pre-amplified cDNA was purified by 0.6x volume of AMpure XP beads, eluted in 14 ul nuclease free water, quantified using the Quant-iT PicoGreen dsDNA Assay Kit and size distributions were checked on a high-sensitivity DNA chip. cDNA was then diluted to 200⍰pg/μl, and 1 ul was used for tagmentation by mixing with 1 μl of tagmentation reaction mixture (10⍰mM TAPS, pH 8.5, 5⍰mM MgCl_2_, 8% PEG8000, 0.08 ul Tn5 mix (illumina)). The reaction was performed at 55°C for 7⍰min, followed by heat inactivation at 80°C for 5⍰min. Library amplification of the tagmented samples was performed using custom-designed index primers and by adding 5⍰μl of PCR mix (1× Phusion High-Fidelity buffer, 0.01⍰unit/ul Phusion High-Fidelity DNA Polymerase, 0.2⍰mM dNTPs, 0.1⍰μM forward indexed primer and 0.1⍰μM reverse indexed primer). Amplification was performed as follows: 3⍰min 72°C; 30⍰s at 95°C; 12 cycles of (10⍰s at 95°C; 30⍰s at 55°C; 30⍰s at 72°C); and 5⍰min at 72°C. Samples from one 96-well plate were pooled, and then purified with 0.6x volume of Ampure XP beads. Libraries were quantified by Qubit™ dsDNA HS Assay Kit and quality checked on a high-sensitivity DNA chip. The size averaged at ~1 Kb should be expected. scRNA-seq libraries were sequenced as 50 bp paired-end reads on the Illumina NovaSeq 6000 platform and importantly, ~20% phiX spike-in was added to resolve the low complexity issue in the first 20 bps. All oligos used are listed in Table S3.

## QUANTIFICATION AND STATISTICAL ANALYSIS

### RNA-seq data analysis

Reads were aligned to the mouse reference genome mm10 using STAR with parameters: --outFilterMismatchNoverLmax 0.2 --outFilterMultimapNmax 1 --outSAMstrandField intronMotif --outSAMmapqUnique 60 --twopassMode Basic --outSJfilterReads Unique --outFilterIntronMotifs RemoveNoncanonical. Gene counts were calculated using featureCounts with parameters: -p -s 2 -t exon, and RefSeq mm10 annotation downloaded from GENCODE. The output gene count tables were used as input into DeSeq2 for normalization and differential expression analysis. For comparing the expression level of different genes within a sample, TPM (transcripts per kilobase million) is calculated as: TPM = (CDS read count * mean read length * 10) / (CDS length * total transcript count)

### ChIP-seq data analysis

Reads were aligned to the mouse reference genome mm10 and dm6 for spike-in samples, using Bowtie2 with default parameters. Reads of quality score less than 30 were removed using samtools and PCR duplicates were removed using picard. Regions in mm10 genome blacklist was removed using bedtools and bigwig files were generated using deeptools and parameters: --binSize 50 --normalizeUsing RPKM --ignoreDuplicates --ignoreForNormalization chrX --extendReads 250 for visualization in IGV. Peaks were called using MACS2 with parameters: -g mm --keep-dup 1 --nomodel --extsize 300. Genomic peak annotation was performed with the R package ChIPseeker considering the region ± 3 kb around the TSS as the promoter. Peak overlapping analysis was performed using the Python package Intervene and visualized using the Python package Matplotlib. Motif discovery was performed using HOMER with default parameters.

For visualization of ChIP-seq, uniquely aligned reads mapping to the mouse genome were normalized using dm6 spike-in as described previously(Orlando et al., 2014). Heatmaps were performed using the functions computeMatrix followed by plotHeatmap and plotProfile from deepTools.

### scRNA-seq data analysis

Raw non-demultiplexed fastq files were processed using zUMIs with STAR to generate expression profiles for both the 5′ ends containing UMIs as well as combined full-length and UMI data. The parameters: find pattern: ATTGCGCAATG, UMI (12-19), cDNA (23-50) were specified for Read 1. The UMI count table containing both intron and exon reads was used by Seurat for downstream analysis. Cells with more than 3000 genes detected and less than 5% of reads mapping to mitochondrial genome were retained. Data were normalized (scTransform) and used for principal component analysis dimensionality reduction, followed by louvain clustering and UMAP dimensionality reduction. Major cell types were readily identifiable by common marker genes: posterior neural progenitor (PNP) expressing Sox3, posterior and ventral neural progenitor (PVNP) expressing Hoxd4, newborn motor neuron (NMN) expressing Neurog2, and motor neuron (MN) expressing Mnx1 and Chat.

Differential gene expression analysis between defined cell clusters was performed using R package presto. The top 500 DEGs were ordered by p value. Averaged expression among the defined cell clusters was scaled by row and used as input to R package ComplexHeatmap for visualization.

## DATA AND CODE AVAILABILITY

The accession numbers for the raw data FASTQ files and processed files for all sequencing data will be available on GEO. Original gel imaging data can be accessed from Mendeley: https://data.mendeley.com/datasets/69vsfxr2n6/draft?a=d996b718-00f6-40c6-9e1b-879175dff621

## ACKNOWLEDGMENTS

We thank Drs. Lynne Vales, Esteban Mazzoni, Anne Schaefer for critical reading of the manuscript as well as past and current Reinberg laboratory members for critical comments and discussions. We also thank the New York University Langone Medical Center (NYULMC) Genome Technology Center for help with sequencing, the NYULMC Cytometry and Cell Sorting Core for help with FACS, and the NYULMC Animal Facility for help with mice housing. This study utilized computing resources at the High-Performance Computing Facility of the Center for Health Informatics and Bioinformatics at the NYULMC. This work was supported by grants to D.R. from the NIH (R01NS100897, R01CA199652) and the Simon Foundation Grant #240344 to D.R.) and the Howard Hughes Medical Institute.

## AUTHOR CONTRIBUTIONS

S.L. and D.R. conceived the project, designed the experiments, and wrote the paper; S.L. performed most of the experiments and all the bioinformatic analysis; K.A.A., C.V.C., M.D. and H.K.M performed the clinical and genetic analyses under the supervision of W.B.D.; S.G.C., I.I., E.E., C.Z., O.Z., M.S., A.S.P., M.S.P., M.T.C., A.W., A.S. and L.G. collected the clinical data; T.K. performed the Nrf1 mouse studies under the supervision of T.C.B. and C.-A.M.; D.R. supervised the study.

