## Supplementary material for "NRF1 Association with AUTS2-Polycomb Mediates Specific Gene Activation in the Brain": Table S1

Supplementary Table 1. *AUTS2* variant (exon 9) and clinical characteristics

|  | **LR15-097** | **LR15-004** | **LR05-007** | **LR15-003** | **LR18-404** | **LR19-314** | **LR19-506** |
| --- | --- | --- | --- | --- | --- | --- | --- |
| cDNA change (NM_015570.2) | c.1483C>T | c.1550C>T | c.1600A>C | c.1603_1626del24 | | | |
| Variant Impact | Nonsense | Missense | Missense | Indel | | | |
| Exon(s) affected (N=19) | 9 | 9 | 9 | 9 | | | |
| Protein change | p.Arg495* | p.Pro517Leu | p.Thr534Pro | p.His535_Thr542del | | | |
| Inheritance | *de novo* | *de novo* | *de novo* | *de novo* | | | |
| Gender | Male | Female | Male | Male | Male | Male | Female |
| Age at exam | 1y | 14y | 15y6mo | 12y | 3y10m | 8y | 15y7mo |
| **Neurodevelopment** |  |  |  |  |  |  |  |
| Intellectual disability | nd | mod-sev | severe | severe | mod-sev | mod-sev | severe |
| Speech-language disability | severe | nd | non-verbal | non-verbal | severe | severe | severe |
| Autism/autistic behavior | 1 | 1 | 1 | 1 | nd | 1 | 1 |
| Hypotonia | 1 | 0 | 1 | 1 | 1 | 1 | 1 |
| Epilepsy | 1 | 1 | 1 | 1 | 0 | 0 | 1 |
| Feeding difficulties (infancy) | 1 | 1 | 1 | 1 | 1 | 1 | 1 |
| **Neuroimaging features** |  |  |  |  |  |  |  |
| Strutural brain anomaly (any) | 0 | 0 | 1 | 1 | 1 | 1 | 0* |
| Corpus callosum hypoplasia | 0 | 0 | 1 | 0 | 1 | 1 | 0* |
| Cerebellar hypoplasia | 0 | 0 | 1 | 0 | 1 | 1 | 0* |
| Posterior fossa (small) | 0 | 0 | 1 | 0 | 1 | 1 | 0* |
| Chiari type 1 | 0 | 0 | 0 | 0 | 1 | 0 | 0* |
| **Dysmorphic features** |  |  |  |  |  |  |  |
| Broad nasal bridge/telecanthus* | 0 | 0 | 1 | 1 | 1 | 1 | 0 |
| Low-set eyebrows | 0 | 0 | 1 | 1 | 1 | 1 | 1 |
| Eyebrows thick | 0 | 1 | 0 | 1 | 0 | 1 | 0 |
| Eyebrows sparse medially | 1 | 0 | 1 | 1 | 1 | nd | nd |
| Nose convex (beaked)* | 0 | 0 | 1 | 1 | 0 | 1 | 0 |
| Prominent nasal tip | 0 | 0 | 1 | 1 | 1 | 1 | nd |
| Low hanging columella* | 0 | 0 | 1 (mild) | 1 (mild) | 0 | 0 | nd |
| Malformed and/ or low-set ears* | 0 | 0 | 1 | 1 | 1 | 0 | 1 |
| Small mouth* | 1 | 0 | 0 | 1 | 1 | 1 | 0 |
| Small jaw* | 1 | 0 | 1 | 0 | 1 | 1 | 1 |
| Clinical diagnosis of RTS suggested | 0 | 0 | 1 | 1 | 0 | 0 | 0 |
| **Skeletal abnormalities** |  |  |  |  |  |  |  |
| Broad phalanges (thumbs/halluces) | 0 | 0 | 1 (mild) | 1 (mild) | 1 (mild) | nd | 0 |
| Kyphosis/scoliosis | 0 | 1 | 1 | 0 | 0 | 0 | 0 |

NOTE:	“1” means + (positive)

“0” means – (negative)

“*” means feature reported in Rubinstein-Taybi syndrome

“nd” means NO DATA
